# RNA reinforces condensate nucleation on chromatin to amplify oncogenic transcription

**DOI:** 10.1101/2025.07.29.665963

**Authors:** Krista A. Budinich, Xinyi Yao, Chujie Gong, Lele Song, Xiaoting Wang, Michelle C. Lee, Kaeli M. Mathias, Qinglan Li, Sylvia Tang, Yiman Liu, Son C. Nguyen, Eric F. Joyce, Yuanyuan Li, Haitao Li, Liling Wan

## Abstract

Aberrant chromatin-associated condensates have emerged as drivers of transcriptional dysregulation in cancer. Although extensive studies have elucidated intrinsic protein sequence features governing their formation, how extrinsic factors within the chromatin environment modulate their assembly and pathogenic function remains poorly understood. Gain-of-function mutations in the histone acetylation reader ENL, found in pediatric leukemia and Wilms tumor, drive oncogenesis by inducing condensate formation at highly selective genomic loci. Here, we uncover a critical role for locally produced gene transcripts in reinforcing the nucleation, chromatin engagement, and oncogenic activity of ENL mutant condensates. Mutant ENL binds RNA in part through a conserved basic patch within its YEATS domain, and this interaction enhances condensate formation both *in vitro* and across diverse cellular contexts. Using a chemically inducible condensate displacement and re-nucleation system, we show that blocking ENL–RNA interactions or transcription impairs condensate reformation at endogenous targets. RNA interactions preferentially enhance mutant ENL occupancy at top-bound, condensate-permissive loci, leading to increased transcriptional bursting and robust gene activation at the single-cell and single-allele level. In mouse models, disrupting ENL–RNA interactions reduces condensate formation and oncogenic transcription in hematopoietic stem and progenitor cells, thereby suppressing ENL mutant-driven leukemogenesis. Together, these findings demonstrate that locally produced RNA transcripts can promote locus-specific nucleation of pathogenic condensates on chromatin, which in turn drive persistent and hyperactive transcription of oncogenic targets and lead to tumorigenesis.

## INTRODUCTION

An increasing number of chromatin-associated proteins have been shown to form nuclear condensates—high local-concentration assemblies often driven by dynamic, multivalent interactions^1^. In some cases, these condensates enrich transcriptional machinery at specific genomic loci to regulate gene expression^2–8^. Dysregulation of condensate formation has been implicated in a range of diseases, notably cancer^9–11^. Genetic mutations or chromosomal translocations in regulatory proteins can induce aberrant condensates, which may be co-opted by cancer cells to activate and sustain oncogenic gene programs^9–11^. Although extensive studies have elucidated intrinsic protein sequence features governing condensate formation, how extrinsic factors within the local chromatin environment regulate their assembly and function remains poorly understood.

RNA has long been recognized as a key component of various biomolecular condensates, including stress granules, the nucleolus, and nuclear speckles^12–15^. Emerging evidence suggests that RNA can also modulate condensates formed by transcriptional regulators. For instance, RNA has been proposed to regulate transcription through a feedback mechanism: low levels of enhancer or promoter RNA produced during transcription initiation promote condensate formation, while the elevated RNA transcripts that accumulate during elongation stimulate condensate dissolution^16^. In pluripotent stem cells, non-coding RNA transcribed from endogenous retroviral elements has been shown to facilitate the hijacking of super-enhancer-associated condensates^17^. Despite these advances, whether and how RNA regulates the formation and function of chromatin-associated pathogenic condensates remains largely unexplored.

To address this question, we leverage oncogenic ENL (Eleven-nineteen-leukemia) mutants found in pediatric acute myeloid leukemia (AML) and Wilms tumor^18–22^. In its wildtype form, ENL binds acetylated histones via its YEATS domain and promotes transcriptional elongation at target genes^23,24^. We previously demonstrated that cancer-associated hotspot mutations in the ENL YEATS domain induce aberrant condensate formation at highly selective target loci, such as the *HOXA* cluster. These condensates are required for ENL mutation-induced hyperactivation of target genes and tumorigenesis^25–27^. While several intrinsic sequence determinants within oncogenic ENL have been shown to mediate condensate formation^26^, how extrinsic factors regulate these condensates remains unclear. Given the role of RNA in other nuclear condensates^12^ and emerging evidence that many transcriptional regulators interact with RNA^28–33^, we hypothesized that RNA may modulate ENL mutant condensates. The robust condensate-forming ability, high locus specificity, and potent oncogenic activity of ENL mutants make them an ideal model for dissecting the regulatory mechanisms of pathogenic condensates.

Here, we uncover a critical role for locally produced RNA transcripts in promoting the nucleation and function of ENL mutant condensates. We show that mutant ENL binds RNA in part through a conserved basic patch within the YEATS domain, and this interaction enhances condensate nucleation and transcriptional bursting at permissive genomic loci. Even partial disruption of ENL:RNA interactions impairs oncogenic gene activation and effectively abolishes tumorigenesis in mouse models. Together, these findings illustrate how oncogenic condensates hijack locally produced RNA transcripts to facilitate locus-specific nucleation and transcriptional output, thereby establishing RNA as a functional cofactor in condensate-mediated gene dysregulation in cancer.

## RESULTS

### ENL binds RNA *in vitro* and in cells in part through a conserved basic patch in the YEATS domain

A previous study showed that AF9, a paralog of ENL, can bind DNA in part through residues in its YEATS domain^34^. To assess whether ENL shares this property, we performed electrophoretic mobility shift assay (EMSA) using purified full-length wildtype ENL (ENL-WT) and the oncogenic ENL-T1 mutant, which harbors the most common of the eight patient-derived hotspot mutations found within ENL’s YEATS domain (Fig. 1a). We found that ENL-WT and ENL-T1 similarly bound double-stranded DNA (dsDNA) in a concentration-dependent manner (Extended Data Fig. 1a, b).

**Fig. 1.**
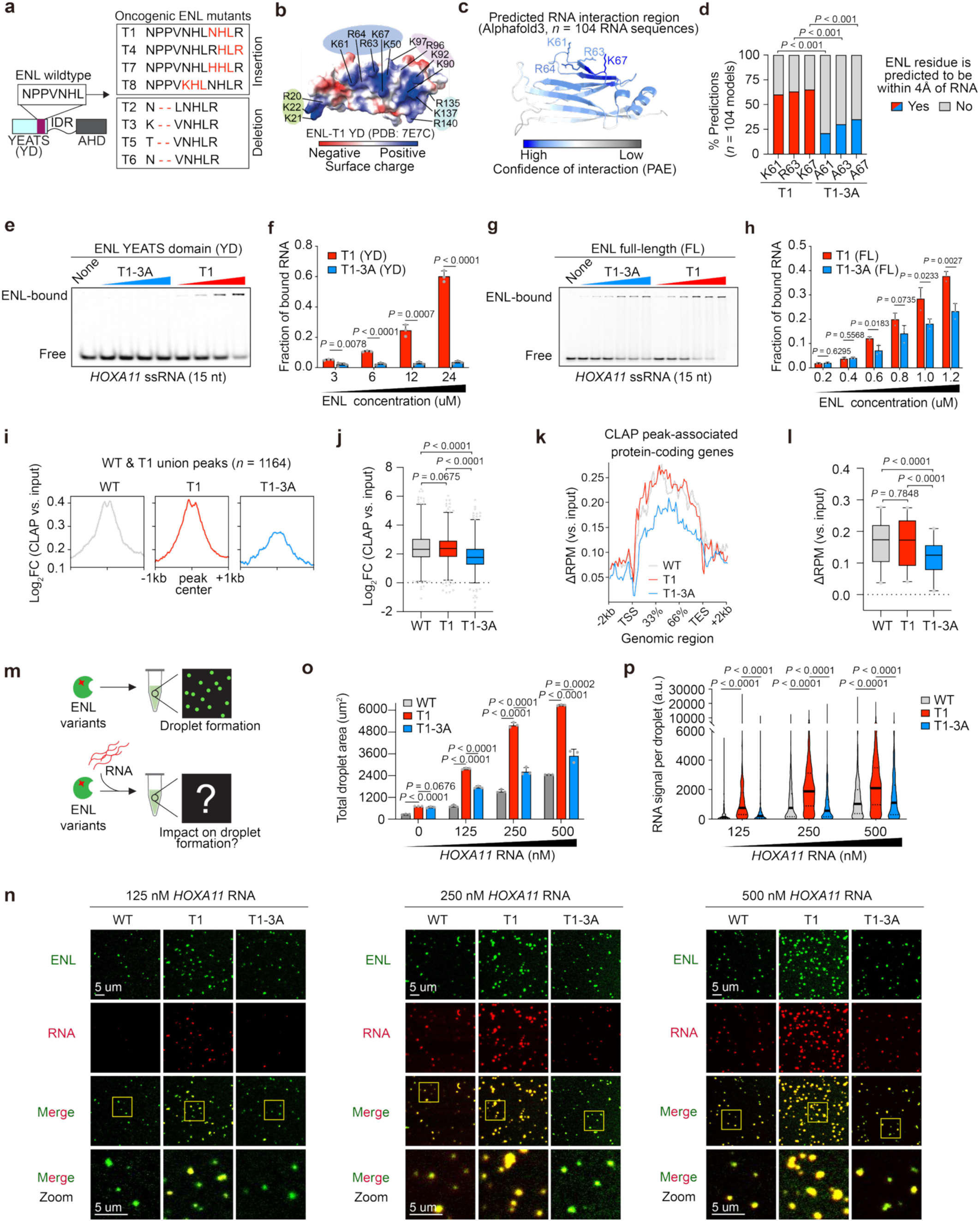
ENL binds RNA in part through a conserved basic patch in the YEATS domain. **a**, Schematic of ENL wildtype protein sequence and its oncogenic hotspot mutations, T1-T8. Mutations are shown in red text. YEATS, Yaf9, Enl, Af9, Taf14, Sas5; IDR, intrinsically disordered region; AHD, ANC1 homology domain. **b,** Surface charge map of ENL-T1 YEATS domain (YD). Residues from positive patches are labeled. **c,** Heatmap depicting confidence scores (PAE, predicted aligned error) of predicted RNA-interacting regions in the ENL-T1 YEATS domain. Predictions generated by Alphafold3. **d,** Percentage of Alphafold3 models in which indicated ENL residue is predicted to be within 4Å of the RNA fragment. *P* values were calculated using two-tailed Fisher’s exact test. **e,** Representative image of EMSA performed with purified ENL YEATS domain protein and a chemically synthesized 15nt *HOXA11* nascent RNA fragment. **f,** EMSA quantification of ENL:RNA binding in (**e**). Data are mean ± s.d. *P* values were calculated using two-tailed unpaired Student’s t test. Representative data shown for *n =* 3 replicates. **g,** Representative image of EMSA performed with purified full-length ENL protein and a chemically synthesized 15nt *HOXA11* nascent RNA fragment. **h,** EMSA quantification of ENL:RNA binding in (**g**). Data are mean ± s.d. *P* values were calculated using two-tailed unpaired Student’s t test. Representative data shown for *n =* 6 biological replicates. **i,** Average RNA signal enrichment at the union list of WT and T1 CLAP peak regions. Values represent Log_2_ fold change of normalized RNA read counts (pulldown vs. input). **j,** Box plots showing Log_2_ fold change of normalized RNA read counts (pulldown vs. input) at WT and T1 union CLAP peaks. Center line indicates median, and box limits are set to 25^th^ and 75^th^ percentiles. *P* values were calculated using two-tailed paired Student’s t test. **k,** RNA signal across gene body regions. Input signals were subtracted from corresponding CLAP pulldowns. RPM, reads per million; TSS, transcription start site; TES, transcription end site. **l,** Box plots showing quantification of (**k**). Center line indicates median, and box limits are set to 25^th^ and 75^th^ percentiles. *P* values were calculated using two-tailed paired Student’s t test. **m**, Schematic of *in vitro* droplet formation assay. **n,** *In vitro* droplet formation by purified recombinant GFP-tagged ENL protein with three concentrations of chemically synthesized and fluorescently labeled nascent *HOXA11* RNA. Yellow box indicates zoom-in region. **o,** Quantification of total droplet area per image field. Data are mean ± s.d. *P* values were calculated by two-tailed unpaired Student’s t test. Representative data shown for *n =* 3 replicates. **p,** Total RNA signal (pixel intensity) per ENL droplet. a.u., arbitrary units. Data are mean ± s.d. *P* values were calculated by two-tailed unpaired Student’s t test. Representative data shown for *n =* 3 replicates.

To test whether ENL can also bind RNA, we performed EMSA with a single-stranded nascent RNA (ssRNA) fragment derived from *HOXA11*—a known ENL mutant target gene whose nascent transcripts are detected within cellular ENL mutant condensates^26^. We found that ENL-WT and ENL-T1 bound *HOXA11* RNA with largely similar affinities (Extended Data Fig. 1c, d). These results indicate that ENL harbors an intrinsic RNA-binding capacity that is preserved in oncogenic ENL mutants. Given the strong enrichment of target RNA transcripts in ENL mutant condensates^26^ and the reported role of RNA in modulating transcriptional condensates^16^, we focused our study on the role of ENL:RNA interactions in condensate formation and function.

We next sought to identify the region within ENL that mediates RNA interaction. Given that AF9 binds DNA through its YEATS domain, we examined the electrostatic surface potential of ENL’s YEATS domain and identified four positively charged surface patches that could engage with negatively charged RNA (Fig. 1b). To evaluate RNA-binding propensity, we modeled interactions between the ENL YEATS domain and ∼100 diverse RNA sequences using AlphaFold3 (Extended Data Fig. 1e and Supplementary Table 1). This analysis highlighted the basic patch encompassing residues K61/R63/R64/K67 as the region most likely to interact with RNA (Fig. 1c). Notably, several positively charged residues within this region align with the DNA-binding residues identified in AF9^34^ (Extended Data Fig. 1f). Structural inspection revealed that among these residues, K61, R63, and K67 are surface-exposed and accessible, whereas R64 is less exposed (Extended Data Fig. 1g).

To test the contribution of this region to RNA binding, we generated a charge-neutralizing triple alanine mutant (K61A/R63A/K67A, hereafter ‘3A’) predicted to preserve the overall YEATS domain structure (Extended Data Fig. 1h). AlphaFold3 modeling showed that the T1-3A mutant exhibited significantly fewer predicted contacts with RNA compared to T1, suggesting a potential role for these residues in RNA interaction (Fig. 1d and Extended Data Fig. 1i). To experimentally test these predicated interactions, we performed EMSA using purified ENL YEATS domains and nascent *HOXA11* RNA fragments of two different lengths. Comparison of ENL-bound RNA fractions revealed that the 3A mutation almost completely abolished the RNA-binding capability of the ENL-T1 YEATS domain (Fig. 1e, f and Extended Data Fig. 1j, k), indicating that this basic region serves as the primary RNA-binding surface within the YEATS domain. We next performed EMSA with full-length ENL proteins using these same *HOXA11* RNA fragments. The 3A mutation consistently reduced RNA binding of full-length ENL-T1, although the effect was partial (Fig. 1g, h and Extended Data Fig. 1l, m). These results suggest that additional regions of ENL may contribute to RNA binding. Given that ENL’s intrinsically disordered region (IDR) contains a concentrated stretch of positively charged residues^26^, we created an ENL-T1 variant in which 21 lysines within this region were substituted with glutamine (IDR(K/Q_21_)). Compared with ENL-T1, this variant exhibited a partial reduction in RNA binding (Extended Data Fig. 1n, o). Collectively these data indicate that ENL binds to RNA through both the YEATS domain and IDR (Extended Data Fig. 1p). However, because large-scale perturbation of the IDR (e.g. K/Q_21_) may also disrupt non-RNA-dependent multivalent interactions required for condensate formation^26^, we focused on the 3A YEATS domain variant as a partial RNA-binding–deficient mutant to directly examine the role of RNA interactions in ENL condensate formation and function.

To determine whether ENL binds RNA in cells, we performed CLAP-seq (Covalent Linkage and Affinity Purification followed by sequencing), an adaptation of CLIP-seq optimized to more directly detect protein–RNA interactions under denaturing conditions^35^. We expressed Halo-tagged ENL-WT and ENL-T1 at near-endogenous levels in HEK293 cells (Extended Data Fig. 2a), UV-crosslinked protein–RNA complexes, affinity-purified ENL and its associated RNAs using Halo ligand resin under denaturing conditions, and sequenced the recovered RNAs (Extended Data Fig. 2b). Comparison of ENL-WT and ENL-T1 CLAP pull-downs with matched inputs identified ∼1200 peaks across both genotypes, with most peaks showing comparable enrichment over input in both WT and T1 conditions (Fig. 1i, j and Extended Data Fig. 2c and Supplementary Table 2). These results indicate that ENL proteins bind RNA in cells and that the oncogenic T1 mutation does not substantially alter the RNA association profiles of ENL. Compared with input, ENL-WT and T1 CLAP peaks were strongly enriched in intronic regions (Extended Data Fig. 2d), suggesting that ENL primarily associates with unspliced pre-mRNA. Moreover, ENL-associated RNAs were predominantly derived from protein-coding genes, consistent with ENL’s known localization to transcribed gene bodies^23,27^ (Extended Data Fig. 2e), and ENL-WT and T1 also exhibited comparable RNA association at these genes (Fig. 1k, l and Extended Data Fig. 2f-h).

To determine whether the 3A mutation affects RNA binding of ENL-T1 in cells, we performed CLAP analysis with cells expressing the ENL-T1-3A transgene. Compared to ENL-WT and ENL-T1, the ENL-T1-3A variant exhibited a partial but significant reduction in RNA association both at the ENL CLAP peaks (Fig. 1i, j and Supplementary Table 2) and across the peak-associated gene bodies (Fig. 1k, l and Extended Data Fig. 2f-h and Supplementary Table 3).

Altogether, our *in vitro* and cellular findings demonstrate that ENL intrinsically binds RNA and that this ability, which is preserved in the oncogenic mutants, depends on a conserved basic patch within its YEATS domain.

### RNA enhances droplet formation of oncogenic ENL mutant *in vitro*

We next asked whether RNA contributes to ENL condensate formation by performing *in vitro* droplet assays using purified full-length ENL protein in the presence or absence of RNA (Fig. 1m). We first tested whether addition of a chemically synthesized nascent RNA fragment of *HOXA11,* a known ENL-T1 target gene, impacts droplet formation. Quantification of total droplet area revealed that ENL-T1 formed droplets more readily than ENL-WT in the absence of RNA (Fig. 1n, o)., consistent with increased self-association conferred by the oncogenic mutation^26^. Addition of RNA further enhanced droplet formation of both ENL-WT and ENL-T1 in a dose-dependent manner, with ENL-T1 showing a more pronounced increase (Fig. 1n, o).

To determine whether this RNA responsiveness depends on the RNA-binding basic patch in the YEATS domain, we tested the T1-3A variant. Importantly, the 3A mutation did not impair the intrinsic droplet-forming abilities conferred by the T1 mutation, as observed in the absence of RNA (Fig. 1n, o). Rather, it specifically reduced RNA-dependent enhancement of droplet formation (Fig. 1n, o). Droplet assays performed with total RNA extracted from whole cells or from the nuclear fraction showed consistent results (Extended Data Fig. 3a-d). Fluorescent labeling of *HOXA11* RNA revealed significantly reduced RNA signal within individual T1-3A droplets compared with T1 droplets (Fig. 1n, p), indicating impaired RNA incorporation into T1-3A condensates. Together, these results suggest that RNA promotes ENL-T1 condensate formation *in vitro* at least in part through interactions with the basic patch of its YEATS domain.

### RNA binding promotes ENL mutant condensate formation in cells

We previously showed that oncogenic ENL mutants form discrete condensates at specific genomic targets when ectopically expressed at near-endogenous levels in both human embryonic kidney-derived HEK293 cells and murine hematopoietic stem and progenitor cells, leading to aberrant target gene activation and leukemogenesis^26,27^. To ensure that condensate formation is not an artifact of overexpression, we focused on the HEK293 system and used a doxycycline-inducible approach to titrate FLAG-tagged ENL-T1 expression (Extended Data Fig. 4a-c). Immunofluorescence (IF) using an anti-FLAG antibody revealed that even when expressed at ∼30% of endogenous ENL levels, ENL-T1, but not ENL-WT, readily formed nuclear condensates and activated target gene expression (Extended Data Fig. 4d-f). Similar results were obtained in *ENL*-knockout HEK293 cells reconstituted with FLAG-tagged ENL-WT or ENL-T1 expressed at near-endogenous levels (Extended Data Fig. 4g–k). Moreover, CRISPR/Cas9-mediated knock-in of the T1 mutation into the endogenous *ENL* locus consistently resulted in condensate formation and target gene induction (Extended Data Fig. 4l-q). Collectively, these results demonstrate that oncogenic ENL mutants potently induce condensate formation and activation of endogenous target genes at physiologically relevant levels, providing a tractable model to dissect how pathogenic condensates are regulated by extrinsic factors such as RNA.

To test whether RNA binding impacts condensate formation in cells, we expressed doxycycline-inducible FLAG-ENL variants at near-endogenous levels in HEK293 cells (Fig. 2a, b). We focused on two of the most common oncogenic mutants: ENL-T1 (insertion) and ENL-T2 (deletion) (Fig. 1a), and generated corresponding RNA-binding-deficient variants (T1-3A and T2-3A). Immunofluorescence using an anti-FLAG antibody revealed that ENL-T1 and -T2 formed discrete nuclear condensates not observed with ENL-WT (Fig. 2c, d). Notably, the 3A mutation partially reduced the number of condensates per cell, indicating that RNA binding promotes the condensate-forming ability of ENL-T1 and ENL-T2 (Fig. 2c, d). To further validate this, we titrated ENL-T1 and ENL-T1-3A expression to levels at or below endogenous ENL protein (30–100% of endogenous levels) (Extended Data Fig. 5a-c). Across this expression range, ENL-T1 formed more condensates per cell than ENL-T1-3A, and even at ∼30% of endogenous levels, ENL-T1 still produced more condensates than ENL-T1-3A expressed at ∼100% of endogenous levels (Extended Data Fig. 5b–e). Similar trends were observed in *ENL*-knockout cells reconstituted with FLAG-ENL variants at physiologically relevant levels, indicating that these effects are not attributable to overexpression (Extended Data Fig. 5f–i).

**Fig. 2.**
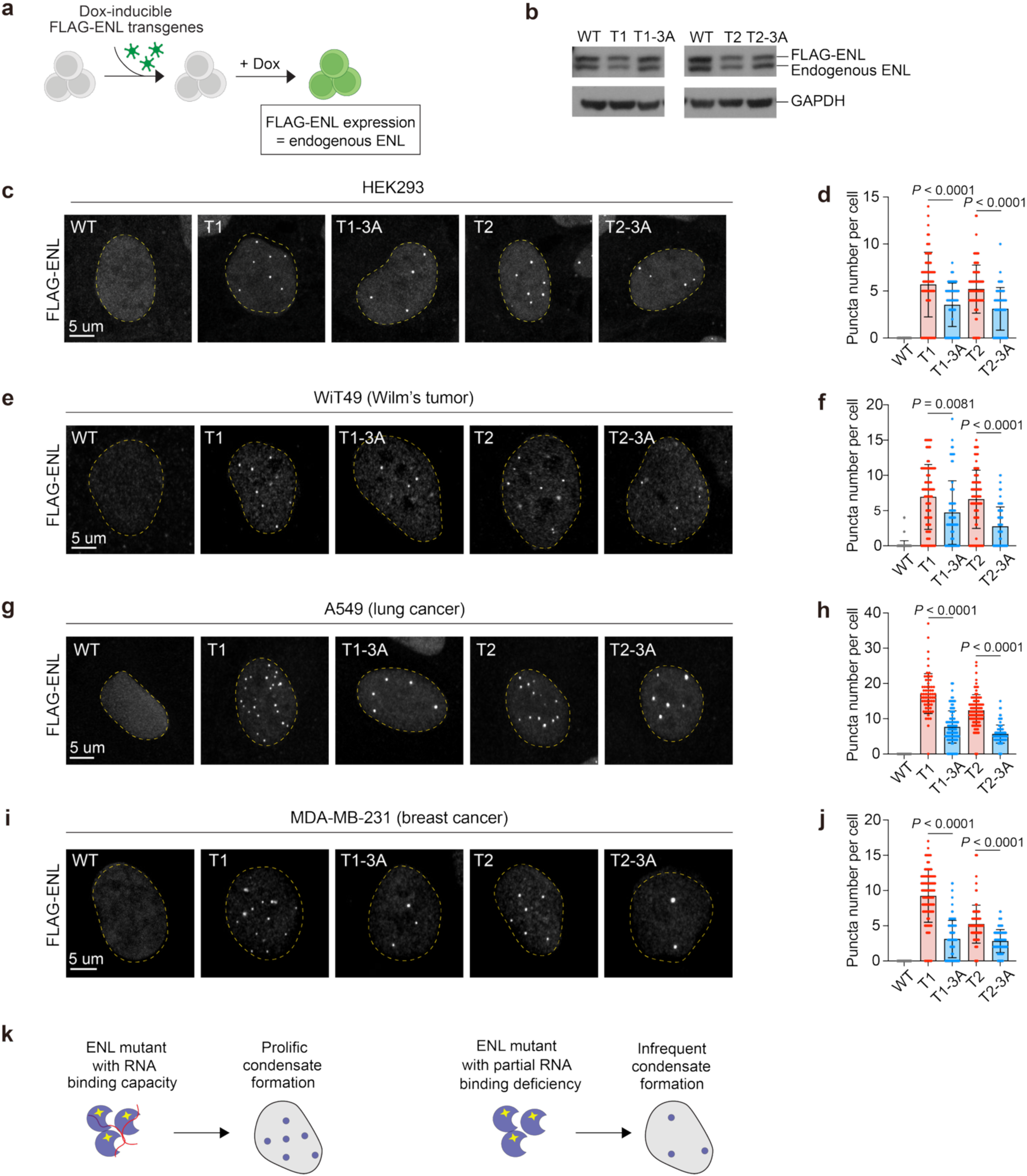
Interactions with RNA promote condensate formation by oncogenic ENL. **a**, Schematic of dox-inducible transgene transduction in cells. **b,** Western blot showing similar expression of ENL transgenes to each other and to endogenous ENL in HEK293 cells. **c, e, g, i,** Representative IF staining of FLAG-ENL in HEK293 (**c**), WiT49 (**e**), A549 (**g**), and MDA-MB-231 (**i**) cells. Yellow dashed line outlines the nucleus. **d, f, h, j,** Quantification of puncta number per cell in HEK293 (**d**), WiT49 (**f**), A549 (**h**), and MDA-MB-231 (**j**) cells. Data are mean ± s.d. *P* values are calculated by two-tailed unpaired Student’s t test. Representative data shown for *n =* 3 replicates. **k,** Schematic depicting summary of results.

In addition to perturbing RNA binding through the charge-neutralizing 3A mutation, we generated a charge-reversal ‘3E’ variant in which the positively charged ‘KRK’ patch (Extended Data Fig. 1f) was replaced with negatively charged glutamates. When expressed at near-endogenous levels in HEK293, the 3E variant, like 3A, exhibited a partial reduction in the number of condensates per cell (Extended Data Fig. 5j-l), further supporting a role for this basic region in promoting condensate formation.

To determine whether these phenotypes are conserved across cellular contexts, we expressed the FLAG-ENL variants in three additional cell lines of diverse tissue origin: WiT49 (Wilms tumor), A549 (lung cancer), and MDA-MB-231(breast cancer) (Extended Data Fig. 5m-o). In all contexts, T1 and T2 formed discrete nuclear condensates, whereas the 3A mutation partially but consistently reduced condensate number (Fig. 2e-j).

To confirm that the reduced condensate formation caused by the 3A mutation reflects impaired RNA binding rather than other perturbations, we examined its effects on additional ENL properties. Cycloheximide chase assays in HEK293 cells expressing different FLAG-ENL variants showed no consistent effects of the 3A mutations on the stability of T1 or T2 proteins (Extended Data Fig. 6a, b). Isothermal titration calorimetry (ITC) of purified YEATS domain variants likewise revealed no significant effect of the 3A mutation on histone acetylation binding (Extended Data Fig. 6c). Moreover, subcellular fractionation followed by immunoblotting revealed that ENL-T1 and ENL-T1-3A exhibited similar subcellular distributions (Extended Data Fig. 6d, e). Finally, immunoprecipitation of FLAG-ENL variants showed no appreciable difference in interactions with CDK9 or ELL2, known ENL-interacting partners within the Super Elongation Complex that are enriched in ENL mutant condensates (Extended Data Fig. 6f-i)^26^. Although additional effects of the 3A mutation cannot be fully excluded, these data, together with results from *in vitro* and cellular RNA-binding assays (Fig. 1), strongly support that the 3A mutation primarily impairs ENL’S RNA-binding ability with minimal impact on other molecular properties. Collectively, our findings demonstrate that RNA binding promotes condensate formation by ENL oncogenic mutants under physiologically relevant expression levels and across diverse cellular contexts (Fig. 2k).

### RNA interactions facilitate ENL mutant condensate formation on and transcriptional bursting at endogenous target loci

To determine whether ENL:RNA interactions contribute to condensate formation at endogenous target genes, we performed concurrent FLAG-ENL immunofluorescence staining and DNA-FISH using probes targeting the *HOXA* or *CBX3* loci, two key targets of oncogenic ENL mutants in kidney cell lines (HEK293 and WiT49) and murine hematopoietic stem/progenitors cells^26,27^. In HEK293 cells expressing ENL-T1 or T2, the majority of *HOXA* and *CBX3* DNA alleles colocalized with ENL mutant puncta (Fig. 3a-h), indicating robust and frequent condensate formation at these targets. In contrast, colocalization was markedly reduced in cells expressing T1-3A or T2-3A variants (Fig. 3a-h), consistent with a lower overall number of nuclear condensates that likely reduces the frequency of condensate association at any given target locus. Corroborating our imaging data, chromatin immunoprecipitation followed by quantitative PCR (ChIP–qPCR) showed that ENL-T1 and T2 exhibited markedly increased occupancy at *HOXA* and *CBX3* relative to ENL-WT, and this enrichment was significantly reduced in T1-3A and T2-3A (Fig. 3i, j). Similar results were seen for T1 versus T1-3E comparisons (Extended Data Fig. 7a).

**Fig. 3.**
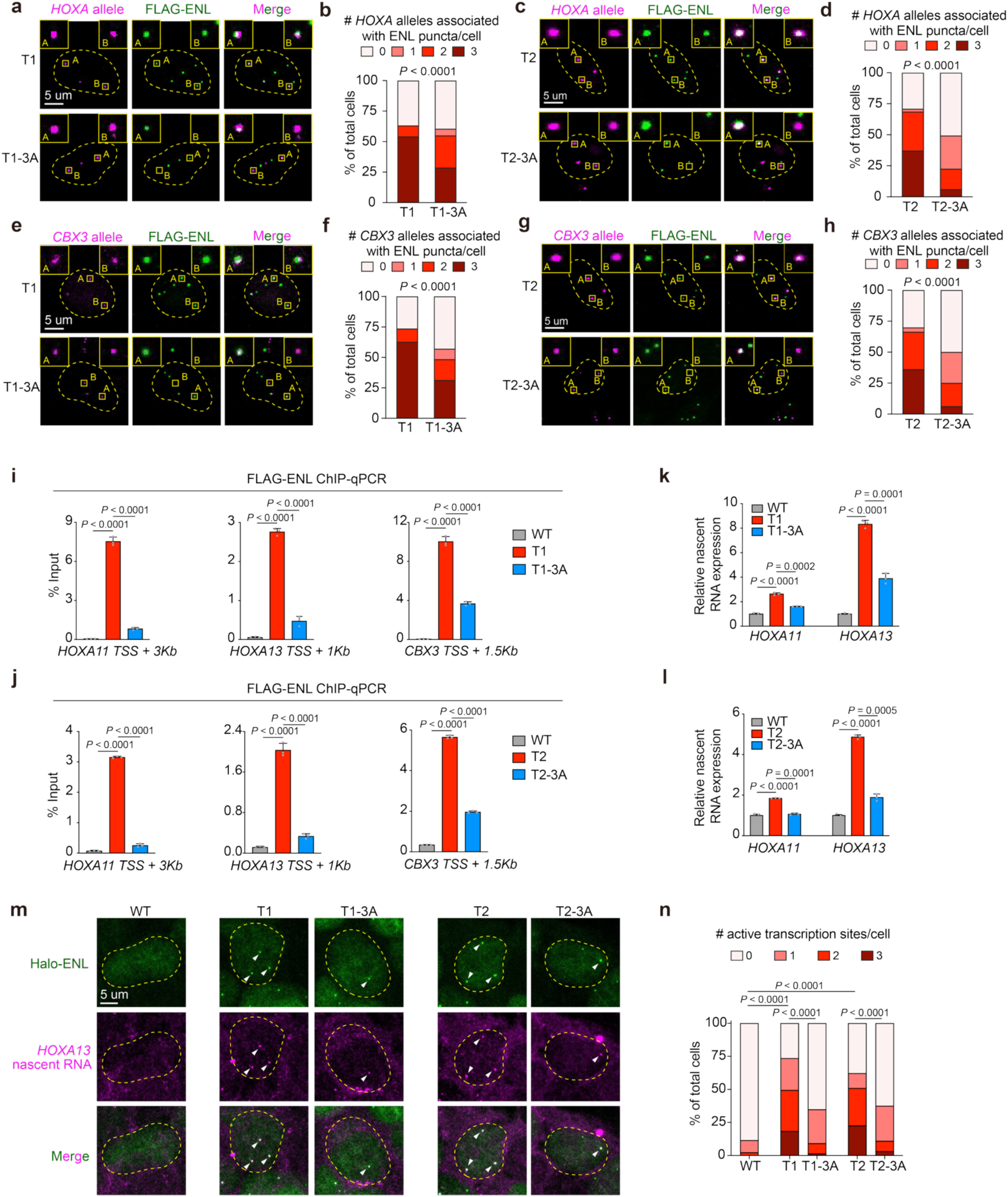
RNA interactions enhance ENL mutant condensate occupancy on and transcriptional bursting of known target loci. **a, c,** Representative *HOXA* DNA-FISH and FLAG-ENL IF staining in HEK293 cells. Yellow dashed line outlines the nucleus **b, d,** Quantification of percentage of cells with indicated number of ENL puncta-associated *HOXA* alleles. *P* values were calculated by two-tailed Fisher’s exact test. Representative data shown for *n =* 3 replicates. **e, g,** Representative *CBX3* DNA-FISH and FLAG-ENL IF staining in HEK293 cells. Yellow dashed line outlines the nucleus **f, h,** Quantification of percentage of cells with indicated number of ENL puncta-associated *CBX3* alleles. *P* values were calculated by two-tailed Fisher’s exact test. Representative data shown for *n =* 3 replicates. **i, j,** ChIP-qPCR analysis of FLAG-ENL variants in HEK293 cells. Data are mean ± s.d. *P* values were calculated by two-tailed unpaired Student’s t test. Triplicate qPCR values are shown. **k**, **l**, RT-qPCR showing nascent *HOXA11* or *HOXA13* expression relative to *GAPDH*. Data are mean ± s.d. *P* values were calculated by two-tailed unpaired Student’s t test. Triplicate qPCR values are shown. **m,** Representative images of nascent RNA-FISH for the *HOXA13* locus and Halo staining for Halo-ENL variants in HEK293 cells. White arrows indicate ENL puncta and RNA-FISH foci. Yellow dashed line outlines the nucleus. **n,** Quantification of percentage of cells with indicated number of nascent RNA-FISH foci. *P* values were calculated by two-tailed Fisher’s exact test. Representative data shown for *n =* 2 replicates.

Given that ENL mutant condensates drive hyperactivation of target genes^26,27^, we next asked whether RNA binding contributes to this function. RT–qPCR for nascent transcripts confirmed reduced expression of key targets in T1-3A, T1-3E, and T2-3A–expressing cells compared to their respective parental mutants T1 and T2 (Fig. 3k, l and Extended Data Fig. 7b). To further demonstrate this transcriptional defect, we titrated ENL-T1 and ENL-T1-3A expression to levels lower than or comparable to endogenous ENL (30-100% of endogenous levels) in HEK293 cells (Extended Data Fig. 5b, c). Similar to condensate formation (Extended Data Fig. 5d, e), the 3A mutation partially reduced transcriptional induction across this expression range, with near-endogenous ENL-T1-3A showing reduced activity even compared to the lowest levels of ENL-T1 expression (Extended Data Fig. 7c).

To further examine transcriptional activity at single-cell and single-allele resolution, we performed nascent RNA-FISH targeting intronic *HOXA13* transcripts and co-stained for Halo-tagged ENL in HEK293 cells. ENL-T1 and ENL-T2 expression led to a significant increase in detected RNA-FISH foci per cell compared to ENL-WT, and these RNA-FISH foci were highly colocalized with ENL-T1 and ENL-T2 condensates (Fig. 3m, n and Extended Data Fig. 7d, e). These results suggest that the presence of ENL mutant condensates likely boosts the transcription bursting frequency at the *HOXA* locus. However, the T1- and T2-induced increases in transcription bursting frequency were largely abrogated by the 3A mutation (Fig. 3m, n). Of note, RNA-FISH foci in T1-3A and T2-3A –expressing cells were still largely occupied by the remaining condensates, further strengthening the link between condensate formation and transcriptional activation (Extended Data Fig. 7d, e).

Together, these results suggest that interactions with RNA facilitate condensate formation by oncogenic ENL at endogenous target loci, thereby increasing transcriptional bursting frequency and gene activation.

### RNA interactions promote re-nucleation of ENL condensates at target loci following acute chemical displacement

As disruption of ENL:RNA interactions via the 3A mutation reduces condensate number (Fig. 2c-j), we hypothesized that RNA binding facilitates condensate nucleation at genomic targets. To directly test this, we established a chemically induced condensate displacement and wash-off system to quantify condensate re-nucleation dynamics at endogenous targets (Fig. 4a). DNA-FISH combined with FLAG-ENL IF showed that ENL-T1 condensates frequently colocalize with the *HOXA* locus in HEK293 cells at basal conditions (Fig. 3a, b and 4a: left panel). To monitor condensate re-nucleation at this site, we first displaced condensates using TDI-11055, a small-molecule inhibitor that competitively blocks the interaction between the ENL YEATS domain and acetylated histones^27,36,37^ (Fig. 4a: middle panel). Following inhibitor wash-off, we tracked the re-formation of condensates at the *HOXA* locus over time in cells expressing either ENL-T1 or the RNA-binding-deficient T1-3A mutant (Fig. 4a: right panel, b).

**Fig. 4.**
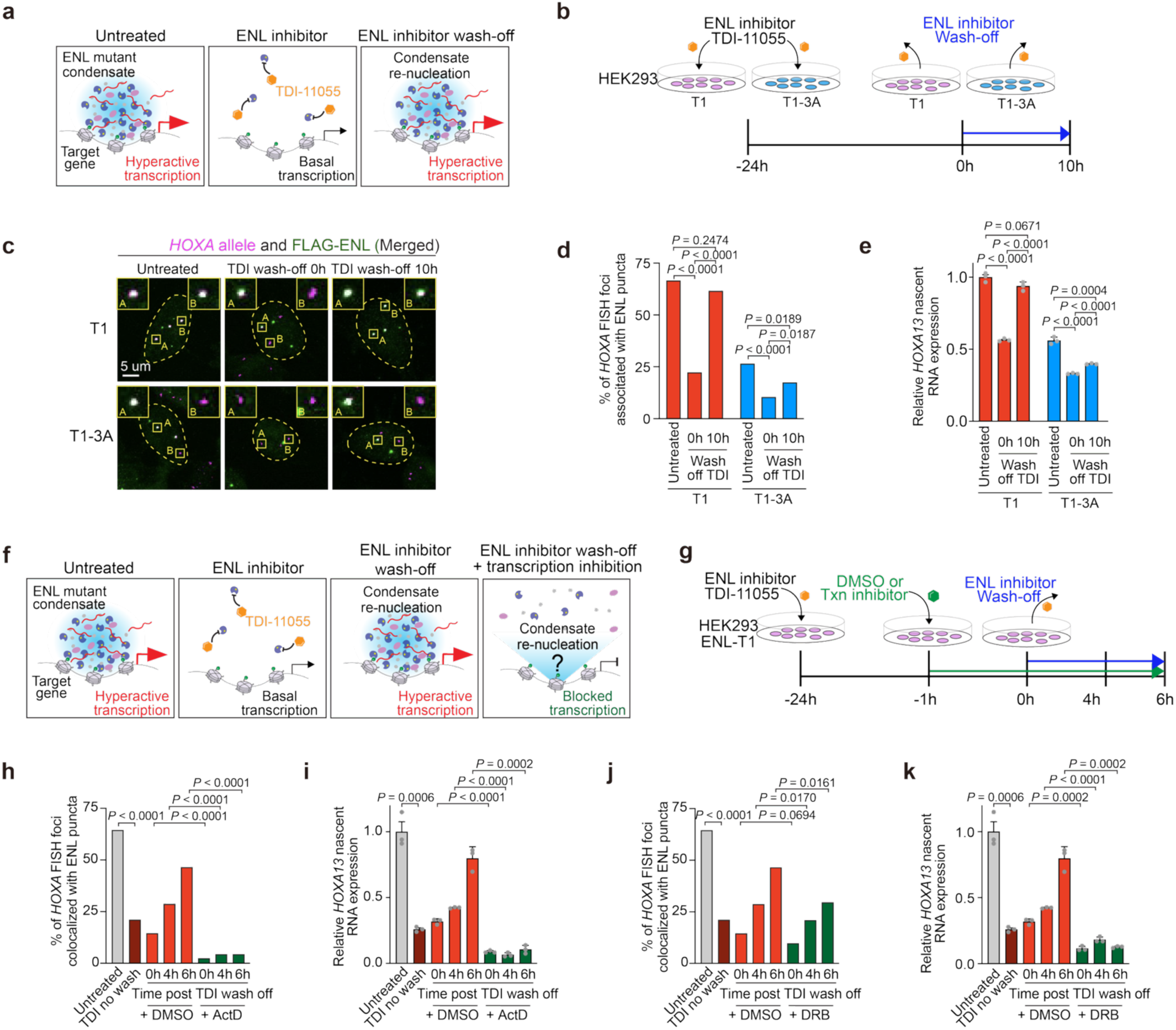
RNA interactions facilitate re-nucleation of ENL mutant condensates at an endogenous target locus after acute chemical displacement. **a**, Schematic depicting experimental design to measure condensate nucleation. Left panel shows ENL mutant condensates reliably occupying target genes. Middle panel shows ENL inhibitor TDI-11055 dislodging ENL condensates from target genes. Right panel shows TDI-11055 wash-off results in the reformation of ENL mutant condensates on target genes. **b,** Schematic of experimental workflow. **c,** Representative *HOXA* DNA-FISH and FLAG-ENL IF staining in HEK293 cells. Yellow dashed line outlines the nucleus. **d,** Quantification of percentage of total *HOXA* alleles associated with ENL mutant puncta in HEK293 cells. TDI, TDI-11055. *P* values were calculated by two-tailed Fisher’s exact test. Representative data shown for *n =* 3 replicates. **e,** RT-qPCR showing nascent *HOXA13* expression relative to *GAPDH*. Data are mean ± s.d. *P* values were calculated by two-tailed unpaired Student’s t test. Triplicate qPCR values are shown. **f,** Schematic depicting experimental design to measure condensate nucleation under transcription inhibition. **g,** Schematic of experimental workflow. **h, j,** Quantification of percentage of total *HOXA* alleles associated with ENL mutant puncta in HEK293 cells. TDI, TDI-11055; ActD, actinomycin D; DRB, 5,6-Dichloro-1-β-D-ribofuranosylbenzimidazole. *P* values were calculated by two-tailed Fisher’s exact test. Representative data shown for *n =* 3 replicates. **i, k,** RT-qPCR showing nascent *HOXA13* expression relative to *18s rRNA*. Data are mean ± s.d. *P* values were calculated by two-tailed unpaired Student’s t test. Representative data shown for *n =* 3 replicates.

In ENL-T1–expressing cells, TDI-11055 treatment markedly reduced the percentage of *HOXA* alleles associated with condensates compared to untreated controls. Upon wash-off, condensate occupancy at *HOXA* loci recovered over a 10-hour period, reaching levels comparable to untreated controls (Fig. 4c, d and Extended Data Fig. 8a). This re-nucleation was accompanied by restored expression of *HOXA11* and *HOXA13* (Fig. 4e and Extended Data Fig. 8b). In contrast, recovery was impaired in T1-3A–expressing cells, both in condensate nucleation and gene activation (Fig. 4c-e and Extended Data Fig. 8a, b).

To further probe of the role of nascent transcripts in condensate nucleation, we applied transcription inhibitors in the wash-off system (Fig. 4f, g). ENL-T1–expressing cells were pre-treated with either DMSO or a transcription inhibitor 1 hour prior to TDI-11055 wash-off. In DMSO-treated cells, condensates rapidly re-formed at the *HOXA* locus, with ∼75% re-nucleation within 6 hours post wash-off (Fig. 4h, j). This re-formation was accompanied by largely restored *HOXA11* and *HOXA13* expression (Fig. 4i, k and Extended Data Fig. 8c, d). In contrast, acute transcription inhibition using Actinomycin D (blocks initiation) or DRB (blocks elongation) significantly impaired condensate re-formation at the *HOXA* locus (Fig. 4h-k and Extended Data Fig. 8c, d).

Together, using a unique chemical strategy to acutely displace and monitor re-formation of condensates at a defined endogenous genomic locus, we show that both RNA-binding capability and ongoing transcription are required for efficient nucleation of ENL-T1 condensates.

### RNA binding preferentially enhances ENL mutant occupancy at condensate-permissive loci

Our DNA-FISH and ChIP-qPCR analyses showed that disrupting ENL:RNA interactions via the 3A mutation reduces chromatin occupancy of oncogenic ENL mutants at condensate-associated genomic loci (Fig. 3a-h). To assess whether RNA binding contributes to ENL mutant occupancy genome-wide or preferentially at specific targets, we performed ChIP-seq for FLAG-ENL variants in HEK293 cells. ENL-T1 and T2 largely occupied the same genomic regions as ENL-WT but with increased occupancy (Extended Data Fig. 9a, b), consistent with increased self-association conferred by these oncogenic mutations^26^. Ranking ENL-T1 or ENL-T2 peaks by their ChIP-seq signal revealed a highly asymmetric distribution, with fewer than 1% of peaks showing disproportionately high occupancy (Fig. 5a and Extended Data Fig. 9c). These top-ranked peaks included validated condensate-associated targets such as the *HOXA* cluster and *CBX3* (Fig. 3a-h). Using the inflection point of the ranked plots, we classified ENL-T1 and ENL-T2 peaks into top- and low-bound groups (Fig. 5a, Extended Data Fig. 9c, and Supplementary Table 4). Top-bound peaks showed a greater fold increase in ENL-T1 or T2 occupancy relative to ENL-WT (Fig. 5b and Extended Data Fig. 9d), suggesting that these target sites are most likely associated with condensates (Fig. 5c).

**Fig. 5.**
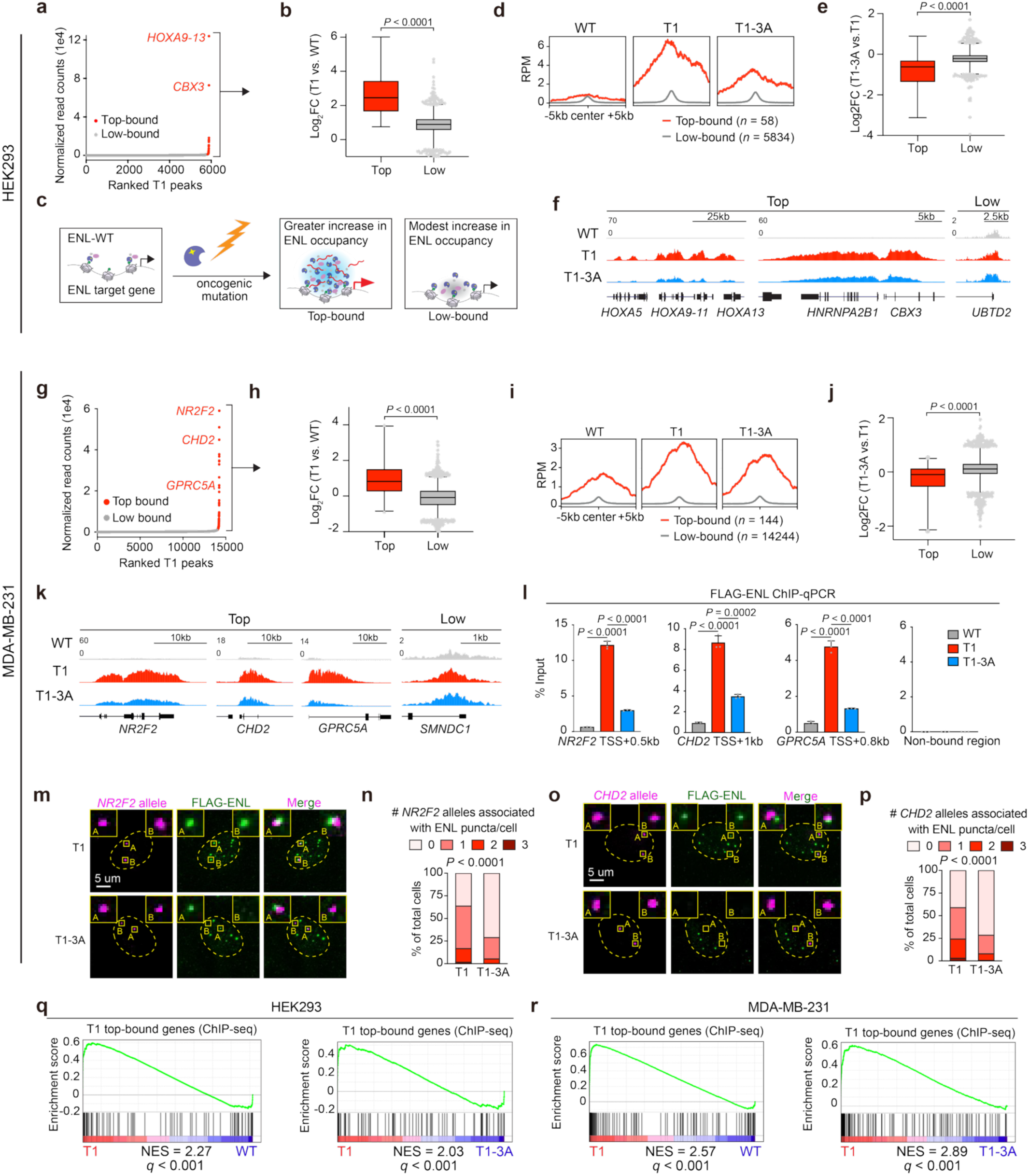
RNA interactions preferentially promote ENL mutant occupancy at condensate-permissive target loci. **a**, ENL-T1 ChIP-seq peaks in HEK293 cells ranked by read count. Top-bound peaks fall to the right of the inflection point and are labeled in red text. Low-bound peaks fall to the left of the inflection point and are labeled in gray. Known condensate-associated targets, *HOXA* and *CBX3*, are indicated. **b,** Box plot showing Log_2_ fold change of T1 vs. WT signal at top or low-bound sites in HEK293. Top, *n =* 58; Low, *n =* 5834. Center line indicates median and box limits are set to 25^th^ and 75^th^ percentiles. *P* values were calculated by two-tailed unpaired Student’s t test. **c,** Schematic showing at most ENL-bound sites (low-bound sites), ENL-T1 modestly gains occupancy compared to ENL-WT. At a very small subset of ENL bound sites (top-bound sites), ENL-T1 exhibits a large gain in chromatin occupancy, likely reflecting gain of condensate formation. **d,** Average chromatin occupancy of ENL at top or low-bound sites. RPM, reads per million. **e,** Box plot showing Log_2_ fold change of T1-3A vs. T1 signal at top or low-bound sites. Top, *n =* 58; Low, *n =* 5834. Center line indicates median and box limits are set to 25^th^ and 75^th^ percentiles. *P* values were calculated by two-tailed unpaired Student’s t test. **f,** Genome browser view of ENL ChIP signal at representative HEK293 top and low-bound genes. **g,** ENL-T1 ChIP-seq peaks in MDA-MB-231 cells ranked by read count. Top-bound peaks fall to the right of the inflection point and are labeled in red text. Low-bound peaks fall to the left of the inflection point and are labeled in gray. **h,** Box plot showing Log_2_ fold change of T1 vs. WT signal at top or low-bound sites in MDA-MB-231. Top, *n =* 144; Low, *n =* 14244. Center line indicates median and box limits are set to 25^th^ and 75^th^ percentiles. *P* values were calculated by two-tailed unpaired Student’s t test. **i,** Average chromatin occupancy of ENL at top or low-bound sites in MDA-MB-231 cells. RPM, reads per million. **j,** Box plot showing Log_2_ fold change of T1-3A vs. T1 signal at top or low-bound sites. Top, *n =* 144; Low, *n =* 14244. Center line indicates median and box limits are set to 25^th^ and 75^th^ percentiles. *P* values were calculated by two-tailed unpaired Student’s t test. **k,** Genome browser view of ENL ChIP signal at representative MDA-MB-231 top and low-bound genes. **l,** ChIP-qPCR of FLAG-ENL variants in MDA-MB-231 cells. Data are mean ± s.d. *P* values were calculated by two-tailed unpaired Student’s t test. Triplicate qPCR values are shown. **m, o**, Representative *NR2F2* (**m**) or *CHD2* (**o**) DNA-FISH and FLAG-ENL IF staining in MDA-MB-231 cells. Yellow dashed line outlines the nucleus. **n, p,** Quantification of percentage of cells with indicated number of ENL puncta-associated *NR2F2* (**n**) or *CHD2* (**p**) alleles. *P* values were calculated by two-tailed Fisher’s exact test. Representative data shown for *n =* 2 replicates. **q, r,** GSEA plots showing that top-bound genes (determined by ChIP-seq) are upregulated in T1 compared to WT or to T1-3A in HEK293 (**q**) and MDA-MB-231 (**r**) cells. NES, normalized enrichment score.

We next asked whether RNA interactions contribute differentially to ENL chromatin occupancy at top-versus low-bound sites. At low-bound sites, ENL-T1 and ENL-T1-3A showed comparable increased occupancy relative to WT (Fig. 5d-f). In contrast, ENL-T1-3A exhibited significantly reduced occupancy relative to T1 at top-bound sites, indicating that RNA binding preferentially reinforces ENL-T1 occupancy at condensate-permissive loci. Similar results were observed for T2 versus T2-3A (Extended Data Fig. 9e-g) as well as for T1 versus T1-3E comparisons (Extended Data Fig. 9h-j).

To test whether this preferential requirement for RNA binding at top-bound sites is conserved across cellular contexts with distinct ENL targets, we performed ChIP–seq in MDA-MB-231 breast cancer cells expressing FLAG-ENL variants at near-endogenous levels (Extended Data Fig. 5o). As in HEK293, ENL-T1 and ENL-T2 exhibited an asymmetric distribution of ChIP–seq signals. However, the top-bound peaks differed substantially between the two cell lines, reflecting cell type-specific targeting (Fig. 5g, Extended Data Fig. 10a, b, and Supplementary Tables 5, 6). Nevertheless, top-bound peaks in MDA-MB-231 also showed a greater increase in occupancy in T1 and T2 relative to WT (Fig. 5h and Extended Data Fig. 10c). Further, the 3A mutation consistently reduced ENL-T1 and T2 occupancy more significantly at top-bound than low-bound sites (Fig. 5i-l and Extended Data Fig. 10d-g).

To determine whether these top-bound sites are indeed associated with condensates in MDA-MB-231 cells, we performed DNA-FISH for two top targets, *NR2F2* and *CHD2*. ENL-T1 and ENL-T2 condensates frequently colocalized with these loci, and this colocalization was reduced in the corresponding 3A variants (Fig. 5m-p and Extended Data Fig. 10h-k).

Together, these findings support a model in which oncogenic mutations amplify ENL occupancy at pre-existing target sites, but only a small, cell type–specific subset is permissive to condensate formation. Although the molecular features that define these loci remain to be elucidated, RNA interactions appear to play a greater role in sustaining ENL occupancy at these sites, likely by facilitating condensate nucleation.

### RNA binding enhances gene activation by oncogenic ENL mutants at condensate-permissive loci

We have shown that ENL mutant condensates increase transcriptional bursting at key condensate-associated targets, and that this effect is reduced by the 3A mutation (Fig. 3m, n). To comprehensively evaluate the contribution of RNA binding to oncogenic ENL mutation-induced transcriptional changes, we performed RNA-seq in HEK293 and MDA-MB-231 cells expressing ENL-T1, ENL-T2, or their corresponding 3A variants at near-endogenous levels (Fig. 2b and Extended Data Fig. 5o). Gene set enrichment analysis (GSEA) using ChIP–seq–defined top-bound genes (Fig. 5a, g, Extended Data Fig. 9c, 10b, and Supplementary Tables 7, 8) revealed upregulation of these targets in ENL-T1 and ENL-T2– expressing cells compared to WT (Fig. 5q, r and Extended Data Fig. 11a, b). This enrichment was attenuated in the T1-3A and T2-3A mutants (Fig. 5q, r and Extended Data Fig. 11a, b). RT–qPCR for nascent transcripts confirmed reduced expression of key targets in T1-3A and T2-3A–expressing cells relative to T1 and T2 in both cell lines (Fig. 3k, l and Extended Data Fig. 11c, d). Together, these results demonstrate that RNA binding enhances the ability of oncogenic ENL mutants to hyperactivate condensate-permissive target genes.

### Partial loss of RNA binding effectively suppresses ENL mutant–driven leukemogenesis

We previously demonstrated that cancer-associated ENL mutants are bona fide drivers of AML using both conditional knock-in mouse models and transplantation models with hematopoietic stem/progenitor cells (HSPCs) expressing ENL mutant transgenes at physiologically relevant levels^27^. Building on this, we sought to determine whether RNA-binding is required for the oncogenic activity of ENL mutants. We isolated HSPC-enriched LSK (Lin⁻Sca1⁺cKit⁺) cells from wildtype C57BL/6 mice and transduced them with FLAG-tagged ENL-WT, T1, or T1-3A transgenes (Fig. 6a). RT–qPCR analysis revealed similar RNA expression levels of the transgenes and the endogenous *Enl* (Extended Data Fig. 12a). Immunofluorescent staining showed that ENL-T1 formed discrete condensates in LSK cells, while ENL-T1-3A exhibited a marked reduction in condensate number per cell (Fig. 6b, c). We next performed DNA-FISH targeting the *Hoxa* cluster and *Meis1*, two key leukemogenic loci previously shown to colocalize with and are activated by ENL mutant condensates in LSK cells^27^. Colocalization of these loci with ENL condensates was significantly more frequent in ENL-T1 than in ENL-T1-3A–expressing cells (Fig. 6d-g), supporting that RNA binding promotes condensate formation at critical oncogenic gene loci in HSPCs.

**Fig. 6.**
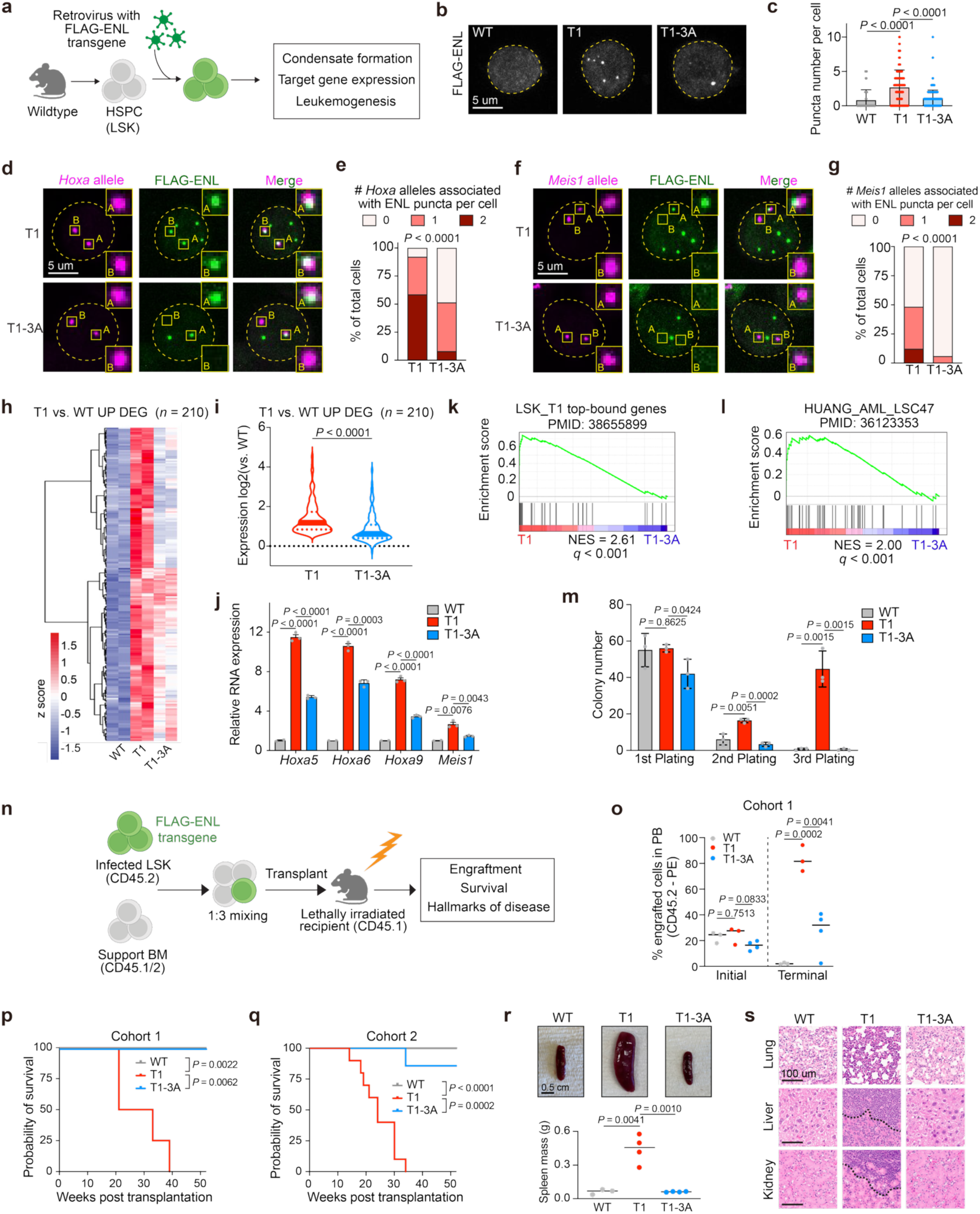
RNA interactions are required for ENL mutant–driven leukemogenesis. **a**, Experimental workflow using hematopoietic stem and progenitor cells (LSK). LSK, Lin^-^Sca^+^cKit^+^. **b,** Representative FLAG-ENL IF staining in LSK cells. Yellow dashed line outlines the nucleus**. c,** Quantification of puncta number per cell. Data are mean ± s.d. *P* values were calculated by two-tailed unpaired Student’s t test. **d, f,** Representative *Hoxa* (**d**) or *Meis1* (**f**) DNA-FISH and FLAG-ENL IF staining in LSK cells. Yellow dashed line outlines the nucleus. **e, g,** Quantification of percentage of cells with indicated number of ENL puncta-associated *Hoxa* (**e**) or *Meis1* (**g**) alleles. *P* values were calculated by two-tailed Fisher’s exact test. **h,** Heatmap depicting z-scores of genes upregulated in T1 vs. WT LSK cells. DEG, differentially expressed genes. **i,** Log2 fold change in expression of T1 vs. WT upregulated genes. Center line indicates median and red or blue dotted lines indicate interquartile range. *P* values were calculated by two-tailed paired Student’s t test. **j,** RT-qPCR showing gene expression of key target loci relative to *Gapdh*. Data are mean ± s.d. *P* values were calculated by two-tailed unpaired Student’s t test. Triplicate qPCR values are shown **k,** GSEA plot showing that ENL-T1 top-bound genes (Extended Data Fig. 9d, n *=* 69) are upregulated in T1 compared to T1-3A in LSK. NES, normalized enrichment score. **l,** GSEA plot showing that genes associated with AML patients (*n =* 47) are upregulated in T1 compared to T1-3A in LSK. NES, normalized enrichment score. **m,** Quantification of colonies formed by LSK expressing indicated ENL variants. Data are mean ± s.d. *P* values were calculated by two-tailed unpaired Student’s t test. Representative data shown for *n =* 3 replicates. **n,** Schematic depicting experimental workflow for bone marrow (BM) transplantation. CD45.1, CD45.2, CD45.1/2 represent leukocyte markers. **o,** Percentage of peripheral blood (PB) composed of cells derived from transplanted LSK at initial vs. terminal timepoints. Each dot represents *n =* 1 mouse. Horizontal line represents median. *P* values were calculated using two-tailed unpaired Student’s t test. **p, q,** Kaplan-Meier survival curves of mice receiving bone marrow transplant across two independent cohorts. *P* value was calculated using Mantel-Cox log-rank test. Cohort 1: WT, *n = 3*; T1, *n =* 4; T1-3A, *n = 4.* Cohort 2: WT, *n = 7*; T1, *n =* 10; T1-3A, *n = 7.* **r,** Representative images (top) and quantification (bottom) of spleen masses harvested from transplanted mice. Each dot represents *n =* 1 spleen. Horizontal line represents median. *P* values were calculated using two-tailed unpaired Student’s t test. **s,** H&E staining. Black dashed line indicates leukemic blast infiltration border.

To assess the transcriptional consequences of disrupting ENL:RNA interactions, we performed RNA-seq in LSK cells expressing ENL-WT, -T1, or -T1-3A. Differential gene expression analysis (fold change > 1.5; *padj* < 0.05) identified five gene clusters (Extended Data Fig. 12b and Supplementary Table 9). Consistent with ENL-T1 functioning as a transcriptional activator, most differentially expressed genes (DEGs) were upregulated from WT to T1 (C3 and C5), while a smaller subset of DEGs were downregulated (C1 and C2) (Extended Data Fig. 12b). Only a limited number of DEGs were uniquely attributable to ENL-T1-3A (C4) (Extended Data Fig. 12b). Focusing on C3 and C5, which represent the majority of DEGs and those upregulated by ENL-T1, we found that the 3A mutation impaired ENL-T1-driven gene activation (Fig. 6h, i), including reduced expression of key condensate-associated targets, the *Hoxa* cluster genes and *Meis1* (Fig. 6j).

Using our previously published ENL-T1 chromatin occupancy dataset generated in LSK cells^27^, we stratified ENL-T1-bound peaks into top- and low-bound groups (Extended Data Fig. 12c and Supplementary Table 10). GSEA revealed that genes associated with top-bound peaks were significantly upregulated in T1 versus WT and remained enriched in T1 compared to T1-3A (Fig. 6k and Extended Fig. 12d). Moreover, genes associated with leukemic stem cells in AML patients were enriched in T1 relative to both WT and T1-3A (Fig. 6l and Extended Data Fig. 12e), indicating that RNA binding contributes to ENL mutant-driven leukemogenic programs.

Finally, we assessed the impact of perturbing ENL:RNA interactions on leukemogenesis. We first performed serial colony formation assays to evaluate self-renewal capacity. ENL-T1–expressing LSK cells maintained robust colony-forming ability over multiple replatings, whereas WT and T1-3A cells rapidly lost this potential (Fig. 6m), indicating that RNA binding is critical for sustaining the T1-induced oncogenic state. We next transplanted LSK cells expressing ENL-WT, T1, or T1-3A into lethally irradiated mice and monitored engraftment over time (Fig. 6n). All groups had similar initial engraftment at two weeks post transplantation (Fig. 6o). However, at the terminal timepoint, defined by maximal disease burden in ENL-T1 mice or by one year post-transplantation for WT and T1-3A, ENL-T1 mice exhibited significantly higher engraftment (Fig. 6o and Extended Data Fig. 12f) and markedly reduced survival across two independent cohorts (Fig. 6p, q). Postmortem analysis confirmed hallmark AML features in ENL-T1 mice, including splenomegaly and leukemic blast infiltration into lung, liver, and kidneys. In contrast, WT and T1-3A mice showed no signs of leukemia even one year post transplantation (Fig. 6r, s). Together, these findings demonstrate that even partial disruption of RNA interactions effectively suppresses leukemogenesis induced by oncogenic ENL mutants, highlighting a critical biological role of RNA in supporting ENL mutant-driven oncogenic activity.

## DISCUSSION

Building on our discovery that ENL directly binds RNA through a basic surface within the YEATS domain, our study establishes an example in which locally produced RNA transcripts reinforce the formation of oncogenic condensates on chromatin, which in turn drives persistent, locus-specific transcriptional activation required for tumorigenesis. Although oncogenic ENL mutations are the primary driver of condensate formation^26^, our genetic and chemical perturbations show that ENL-RNA interactions facilitate condensate nucleation at endogenous genomic targets, thereby promoting local enrichment of mutant ENL and possibly other cofactors that amplify transcriptional bursting at condensate-permissive sites (Fig. 7a). Notably, even partial disruption of RNA interactions through the 3A mutation consistently reduces condensate formation, chromatin occupancy, and target gene activation by oncogenic ENL—and, more strikingly, almost completely abolishes leukemogenesis. These findings suggest that ENL mutant–driven tumorigenesis depends on a critical threshold of oncogenic gene expression, and that RNA interactions help achieve this by reinforcing condensate nucleation and transcriptional activation.

**Fig. 7.**
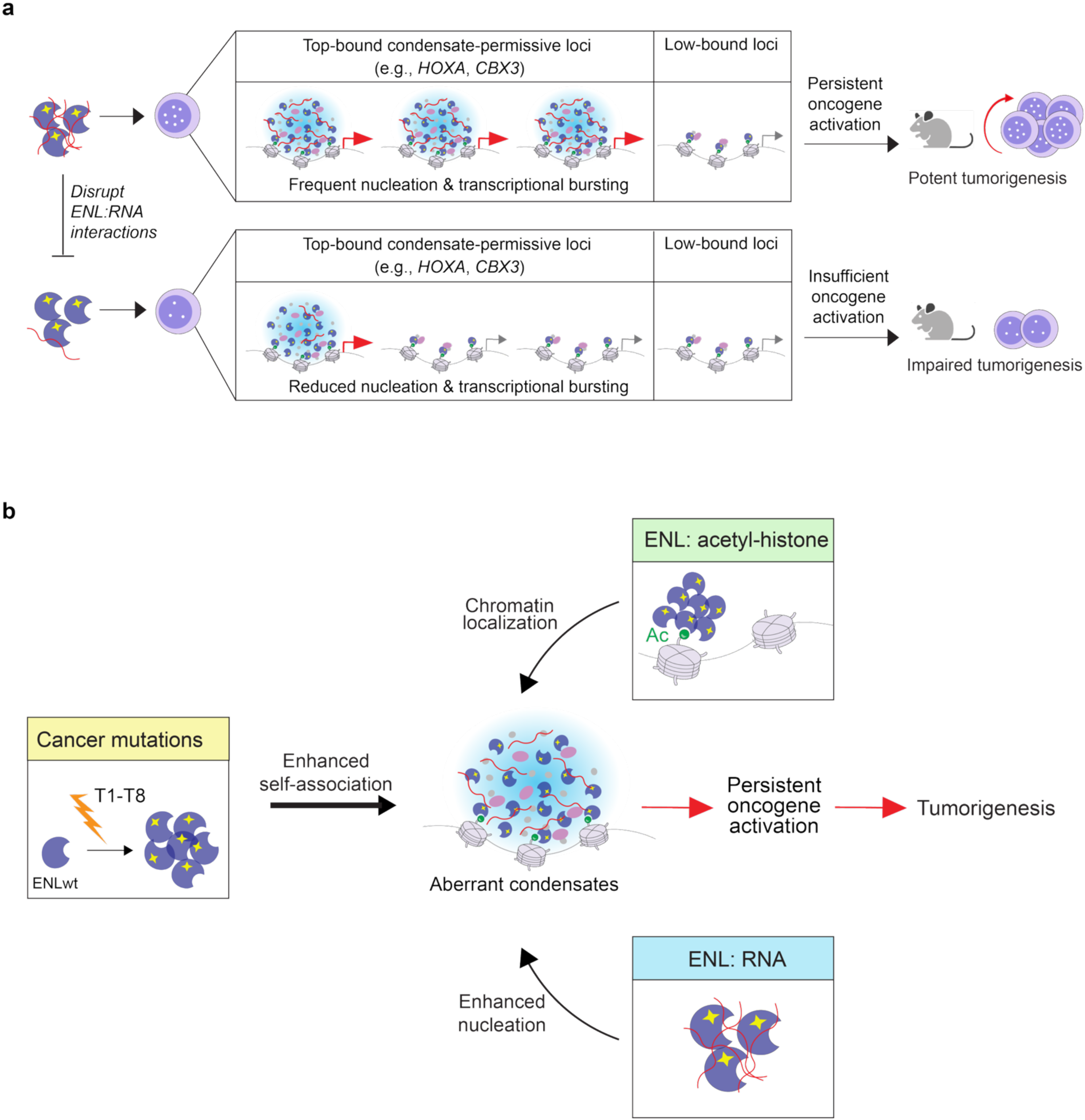
Pathogenic ENL mutant condensates are regulated by complementary mechanisms. **a**, Model illustrating that (Top) oncogenic ENL’s interactions with locally produced RNA transcripts facilitate condensate nucleation at key target genes, thereby promoting frequent transcriptional bursting and persistent activation at top-bound, condensate-permissive loci, ultimately leading to potent leukemogenesis in mouse models. (Bottom) Disruption of these ENL:RNA interactions reduces condensate formation, transcriptional bursting, and target gene activation, thereby impairing tumorigenesis. **b,** Schematic summarizing the multi-layered mechanisms underlying ENL mutant condensate formation and function. Cancer-associated mutations enhance ENL self-association and trigger condensate formation, while intrinsic ENL–acetyl-histone and ENL–RNA interactions facilitate proper chromatin localization and efficient condensate nucleation, respectively. Together, these mechanisms enable robust condensate formation and sustained gene activation at key target loci, thereby achieving the critical threshold of oncogenic gene expression required for tumorigenesis.

Together with our previous work, this study delineates the multifaceted mechanisms underlying ENL mutant condensate formation and function. Cancer-associated mutations (T1-T8) promote ENL self-association and serve as the initiating driver of condensate formation, whereas ENL’s intrinsic binding to histone acetylation is required for condensate association with chromatin ^23,25–27,38^. Here, we identify a third layer of regulation—interactions with locally produced transcripts—that facilitates condensate nucleation. These homotypic and heterotypic molecular interactions cooperate to establish robust condensate formation at specific genomic loci, resulting in sustained hyperactivation of oncogenic targets and potent tumorigenesis (Fig. 7b). Such interplay among oncogenic proteins, chromatin, and local gene transcripts likely represents a general principle underlying the formation and function of other chromatin-associated condensates.

While the involvement of RNA in pathogenic condensates is increasingly recognized, prior studies have primarily focused on how dysregulation of an RNA’s folding, expression, localization, or modification triggers pathogenic condensate formation or alters condensate properties and/or function^12–14^. Our study highlights that condensate-forming oncogenic proteins can co-opt RNA molecules to enhance their condensate nucleation and function. Moreover, whereas noncoding RNAs have long been implicated in pathogenic condensate assembly, our findings suggest that intron-containing RNA transcripts proximal to target genomic loci likely represents the key RNA species that promote ENL mutant condensate nucleation, thus adding local gene transcripts to the growing repertoire of RNA species that contribute to pathogenic condensate biology^12–14^.

The role of RNA identified in ENL mutant condensate formation conceptually aligns with models proposing that basal levels of promoter- or enhancer-derived RNA produced during transcription initiation can promote transcriptional condensate formation. However, our findings suggest that the consequences of RNA interactions in physiological or pathological contexts likely diverge in response to persistent transcriptional bursting. In physiological settings, such as shown with Mediator condensates in mouse embryonic stem cells, rising levels of genic RNA during transcriptional bursts are proposed to disrupt condensates by altering local electrostatic balance, thereby providing a feedback mechanism that fine-tunes gene expression required for development and homeostasis^16^. In contrast, we observe stable condensate formation by oncogenic ENL mutants at key target genes despite exceedingly high transcriptional output driven by persistent bursting. We propose that basal levels of locally produced RNA transcripts engage ENL mutants through multivalent interactions, therefore lowering the energetic barrier for condensate nucleation. Once formed, the high local concentration of ENL mutant proteins within the condensates may buffer charge imbalance caused by increasing RNA levels through direct ENL–RNA interactions, thereby preventing RNA-mediated dissolution, maintaining condensate stability, and sustaining persistent oncogenic transcription required for tumorigenesis.

Beyond mutant ENL, other oncogenic drivers, such as NUP98-HOXA9, EWS-FLI1, and mutant nucleophosmin (NPM1), also form chromatin-associated condensates enriched in transcription regulatory factors^6,39–43^. These proteins often contain intrinsically disordered regions with charge blocks that may be involved in RNA interactions^39,44,45^. Thus, RNA-dependent condensate regulation may represent a broader strategy through which cancer cells achieve persistent pathogenic gene activation, a hypothesis warranting further investigation.

## DATA AVAILABILITY

All CLAP-seq, ChIP-seq, and RNA-seq raw and processed data have been deposited in the Gene Expression Omnibus database under accession numbers GSE300418, GSE300420, GSE300421. All other raw data generated or analyzed during this study are included in this paper, the Extended Data Figures, and the Supplementary Information.

## ACKNOWLEDGEMENTS

We thank members of the Wan Laboratory and the Li Laboratory for scientific input throughout the study; the CDB Microscopy Core at the University of Pennsylvania; the Flow Cytometry and Cell Sorting Core at the Children’s Hospital of Pennsylvania; the Pathology Core at the Children’s Hospital of Pennsylvania; the AFCRI Research Facilities Team at the University of Pennsylvania, and the University Laboratory Animal Resources team at the University of Pennsylvania. The research was supported by funding from the University of Pennsylvania (to L.W.), Pew-Stewart Scholar Award (to L.W.), American Cancer Society Award (to L.W.), Blood Cancer United (formerly The Leukemia & Lymphoma Society) Scholar Award (to L.W.), the National Key Research and Development Program of China (2022YFA1304800, to Y.Li; 2021YFA1300103, 2020YFA0803303, to H.L.), and the National Natural Science Foundation of China (T2488301, 32230019, to H.L.).

## AUTHOR CONTRIBUTIONS

K.A.B, Y.Li, H.L., and L.W. conceived and designed the study. K.A.B performed the cellular, molecular, genomics, and *in vivo* animal studies. X.Y., X.W. and Y.Li performed *in vitro* biochemical studies. C.G. performed bioinformatic analysis of genomics data. Q.L. provided bioinformatics analysis pipelines. M.C.L., S.T., Y.Liu, L.S., and K.M.M. provided technical assistance. S.C.N. and E.F.J. designed and synthesized FISH probes. K.A.B. and L.W. wrote the manuscript with input from H.L., Y.Li, X.Y., L.S., and C.G. H.L. and Y.Li jointly supervised the *in vitro* biochemical studies. L.W. supervised the overall study.

---

**Extended Data Fig. 1.**
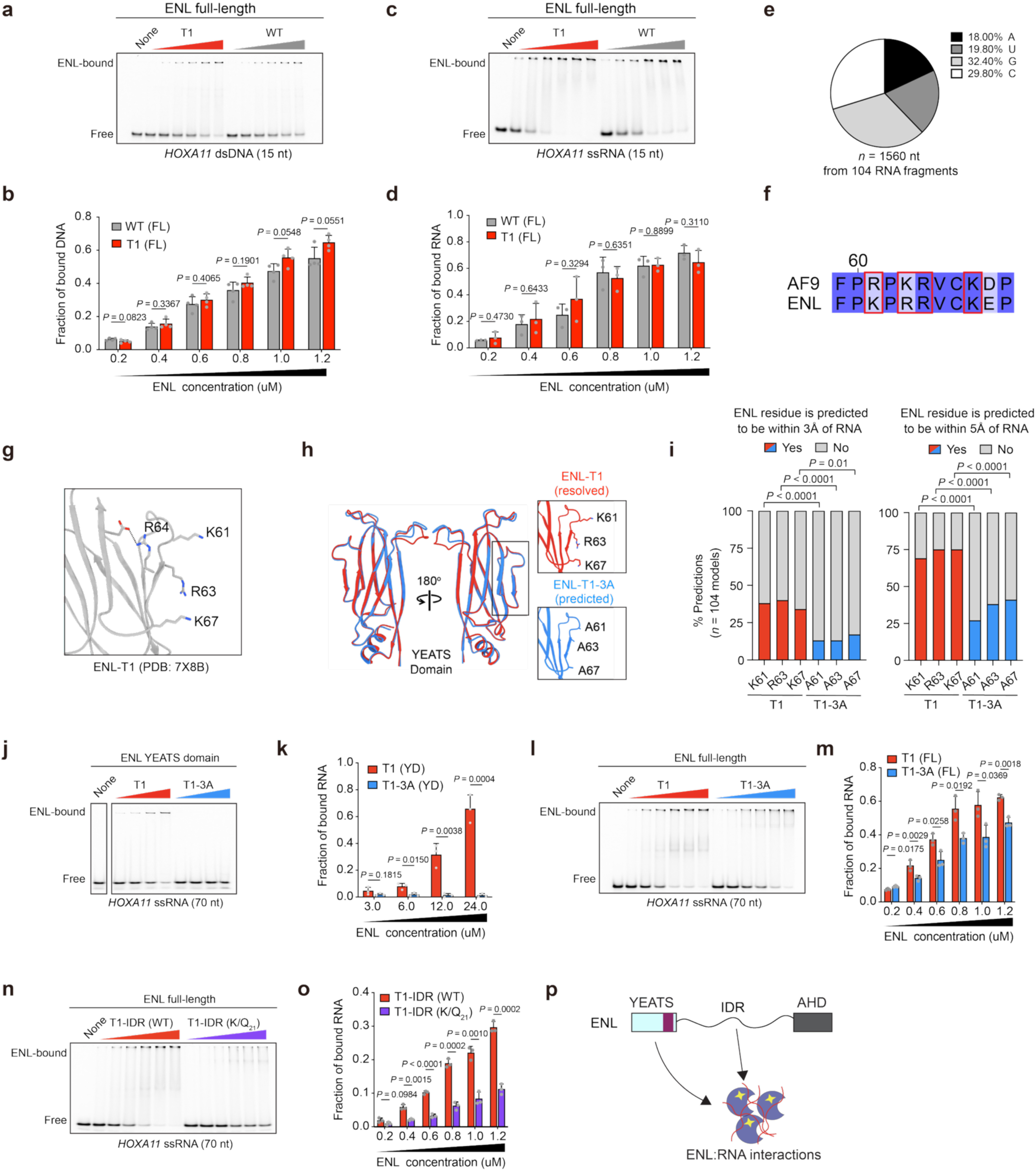
ENL binds RNA in part through a conserved basic patch in the YEATS domain *in vitro*. **a, b,** Representative image (**a**) and quantification (**b**) of EMSA performed with purified full-length ENL protein and chemically synthesized 15nt *HOXA11* dsDNA fragment. Data are mean ± s.d. *P* values were calculated using two-tailed unpaired Student’s t test. Representative data shown for *n =* 3 replicates. **c, d,** Representative image (**c**) and quantification (**d**) of EMSA performed with purified full-length ENL protein and chemically synthesized 15nt *HOXA11* ssRNA fragment. Data are mean ± s.d. *P* values were calculated using two-tailed unpaired Student’s t test. Representative data shown for *n =* 3 replicates. **e,** Distribution of nucleotides in 104 RNA sequences used for Alphafold3 modelling. **f,** Sequence alignment of ENL and ENL paralog, AF9. AF9’s DNA-binding residues are outlined in red. **g,** ENL-T1 structure shows K61, R63, and K67 are surface-exposed, while R64 is less accessible. **h,** Overlay of resolved ENL-T1 and predicted ENL-T1-3A YEATS domain structures. T1-3A structure predicted with Alphafold3. **i,** Percentage of Alphafold3 models in which indicated ENL residue is predicted to be within 3Å or 5Å of the RNA molecule. *P* values were calculated using two-tailed Fisher’s exact test. **j, k,** Representative image (**j**) and quantification (**k**) of EMSA performed with purified ENL YEATS domain protein and *in vitro* transcribed 70nt *HOXA11* ssRNA fragment. Singular gel is shown and was cropped for clarity. Data are mean ± s.d. *P* values were calculated using two-tailed unpaired Student’s t test. Representative data shown for *n =* 3 replicates. **l, m,** Representative image (**l**) and quantification (**m**) of EMSA performed with purified full-length ENL protein and *in vitro* transcribed 70nt *HOXA11* ssRNA fragment. Data are mean ± s.d. *P* values were calculated using two-tailed unpaired Student’s t test. Representative data shown for *n =* 3 replicates. **n, o,** Representative image (**n**) and quantification (**o**) of EMSA performed with purified full-length ENL protein and *in vitro* transcribed 70nt *HOXA11* ssRNA fragment. IDR, intrinsically disordered region. IDR (K/Q_21_), 21 lysines mutated to glutamine. Data are mean ± s.d. *P* values were calculated using two-tailed unpaired Student’s t test. Representative data shown for *n =* 3 replicates. **p,** The YEATS domain and IDR both contribute to ENL RNA binding.

**Extended Data Fig. 2.**
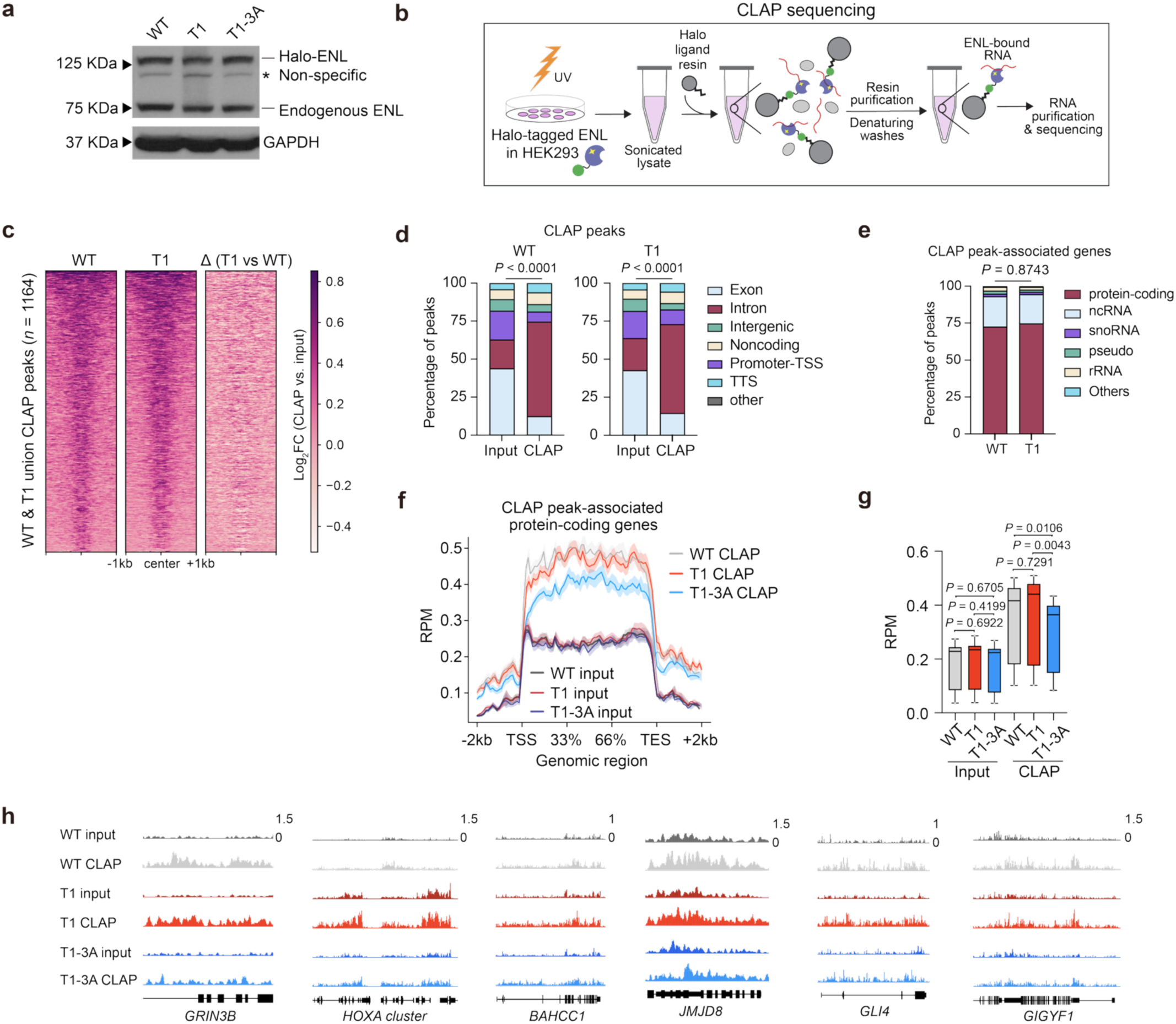
ENL binds RNA in part through a conserved basic patch in the YEATS domain in cells. **a**, Western blot showing comparable expression of ENL transgenes to each other and to endogenous ENL in HEK293 cells. **b,** Schematic of CLAP-sequencing workflow. **c,** Heatmaps showing Log_2_ fold changes of RNA peak signal (CLAP vs. input) on WT & T1 union CLAP peaks. **d,** Distribution of input or CLAP pull-down RNA peaks across genomic regions. TSS, transcription start site; TTS, transcription termination site. *P* values were calculated by two-tailed Fisher’s exact test. **e,** Distribution of genes associated with ENL CLAP pull-down peaks. ncRNA, noncoding RNA; snoRNA, small nucleolar RNA; rRNA, ribosomal RNA. **f**, ENL-bound RNA signal across gene body regions. RPM, reads per million; TSS, transcription start site; TES, transcription end site. **g**, Box plots showing RNA signal (CLAP pull-downs vs. inputs) of peak-associated genes. Center line indicates median, and box limits are set to 25^th^ and 75^th^ percentiles. *P* values were calculated using two-tailed paired Student’s t test. **h**, Genome browser view of ENL-bound RNA signal in inputs and CLAP pull-downs.

**Extended Data Fig. 3.**
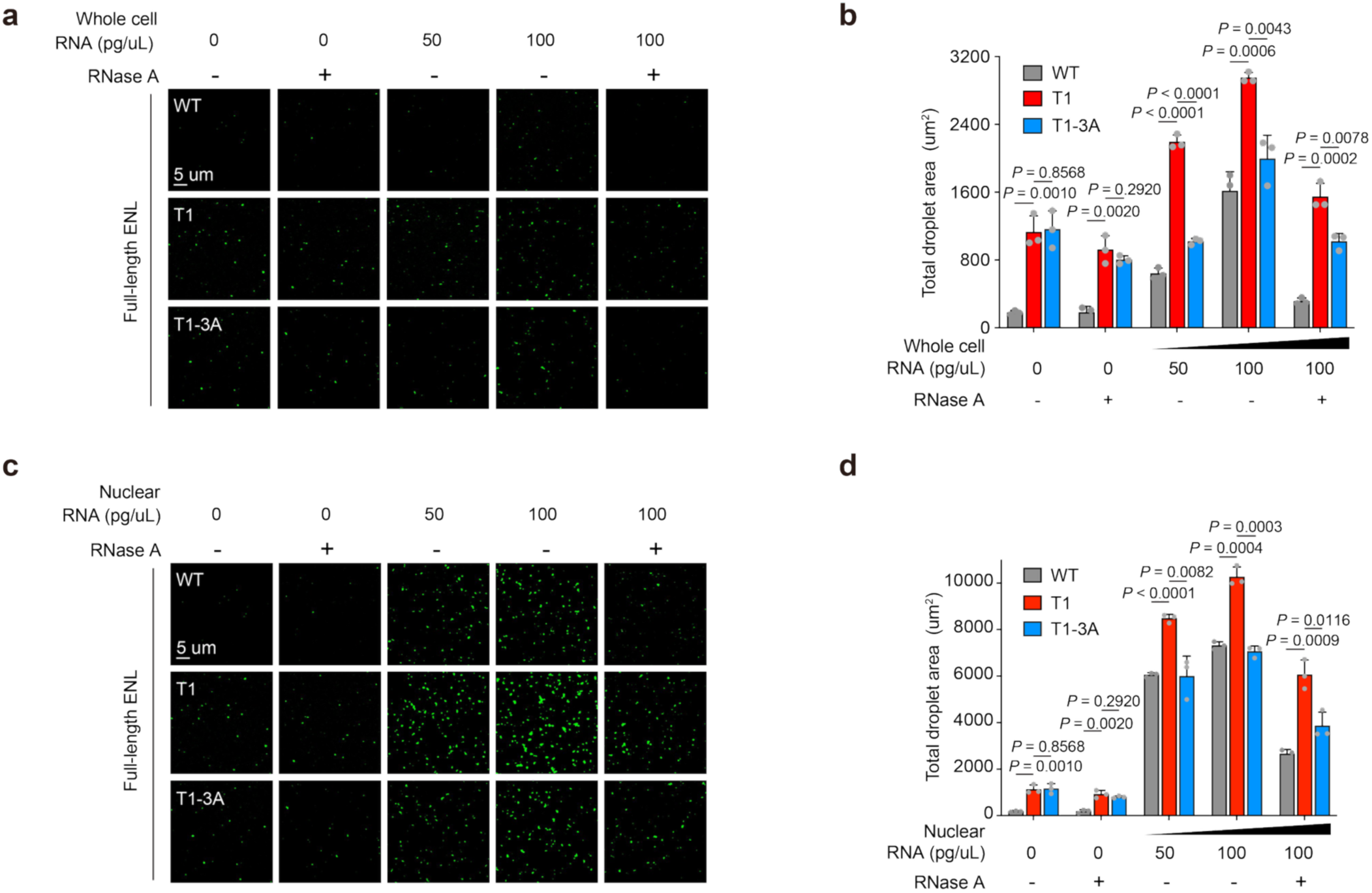
RNA interactions enhance droplet formation of purified ENL protein. **a**, *In vitro* droplet formation by purified recombinant GFP-tagged ENL protein with RNA extracted from whole cell lysate. **b,** Quantification of total droplet area per image field. Data are mean ± s.d. *P* values were calculated by two-tailed unpaired Student’s t test. Representative data shown for *n =* 3 replicates. **c,** *In vitro* droplet formation by purified recombinant GFP-tagged ENL protein with RNA extracted from cell nuclei. **d,** Quantification of total droplet area per image field. Data are mean ± s.d. *P* values were calculated by two-tailed unpaired Student’s t test. Representative data shown for *n =* 3 replicates.

**Extended Data Fig. 4.**
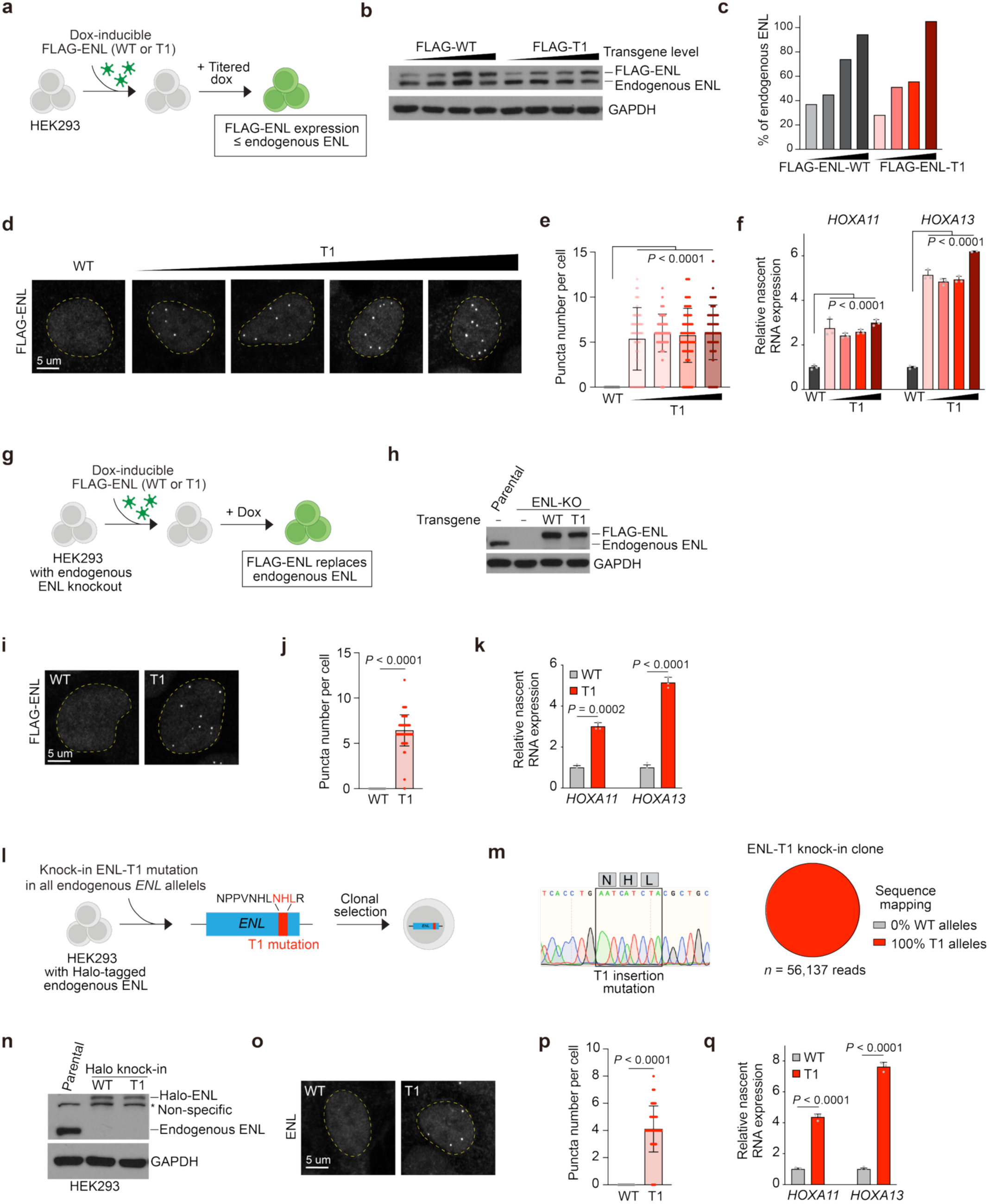
Oncogenic ENL mutants induce aberrant condensate formation and target gene activation at physiologically relevant expression levels in HEK293 cells. **a**, Schematic of dox-inducible transgene transduction in cells. FLAG-ENL variants are expressed at lower than or close to endogenous levels of ENL. **b**, Western blot showing low to near-endogenous expression of FLAG-ENL variants relative to endogenous ENL. **c,** Quantification of (**b)**. FLAG-ENL intensity shown as a percentage of endogenous ENL intensity. **d**, Representative IF staining of FLAG-ENL variants in HEK293 over increasing T1 expression levels. FLAG-WT = ∼endogenous ENL. **e,** Quantification of puncta number per cell. FLAG-WT = ∼endogenous ENL Data are mean ± s.d. *P* values are calculated by two-tailed unpaired Student’s t test. Representative data shown for *n =* 2 replicates. **f**, RT-qPCR showing nascent *HOXA11* or *HOXA13* expression relative to *18s rRNA*. FLAG-WT = ∼endogenous ENL. Data are mean ± s.d. *P* values were calculated by two-tailed unpaired Student’s t test. Triplicate qPCR values are shown. **g,** Schematic of dox-inducible transgene transduction in HEK293 cells with endogenous ENL knockout. FLAG-ENL variants are expressed at near endogenous levels of ENL. **h**, Western blot showing near-endogenous reconstitution of FLAG-ENL variants in ENL-knockout cells. **i**, Representative IF staining of FLAG-ENL variants in ENL-knockout cells. **j,** Quantification of puncta number per cell. Data are mean ± s.d. *P* values are calculated by two-tailed unpaired Student’s t test. Representative data shown for *n =* 2 replicates. **k**, RT-qPCR showing nascent *HOXA11* or *HOXA13* expression relative to *18s rRNA*. Data are mean ± s.d. *P* values were calculated by two-tailed unpaired Student’s t test. Triplicate qPCR values are shown. **l,** Schematic depicting knock-in of T1 mutation into endogenous *ENL* alleles and subsequent clonal selection. **m**, Sanger sequencing trace shows knock-in of T1 mutation into *ENL* exon 4 and CRISPR sequencing confirms knock-in into all *ENL* alleles. **n,** Western blot shows T1-knockin does not impact protein integrity. **o,** Representative IF staining of ENL. Yellow dashed line outlines the nucleus. **p,** Quantification of puncta number per cell. Data are mean ± s.d. *P* values are calculated by two-tailed unpaired Student’s t test. Representative data shown for *n =* 2 replicates. **q**, RT-qPCR showing nascent *HOXA11* or *HOXA13* expression relative to *18s rRNA*. Data are mean ± s.d. *P* values were calculated by two-tailed unpaired Student’s t test. Triplicate qPCR values are shown.

**Extended Data Fig. 5.**
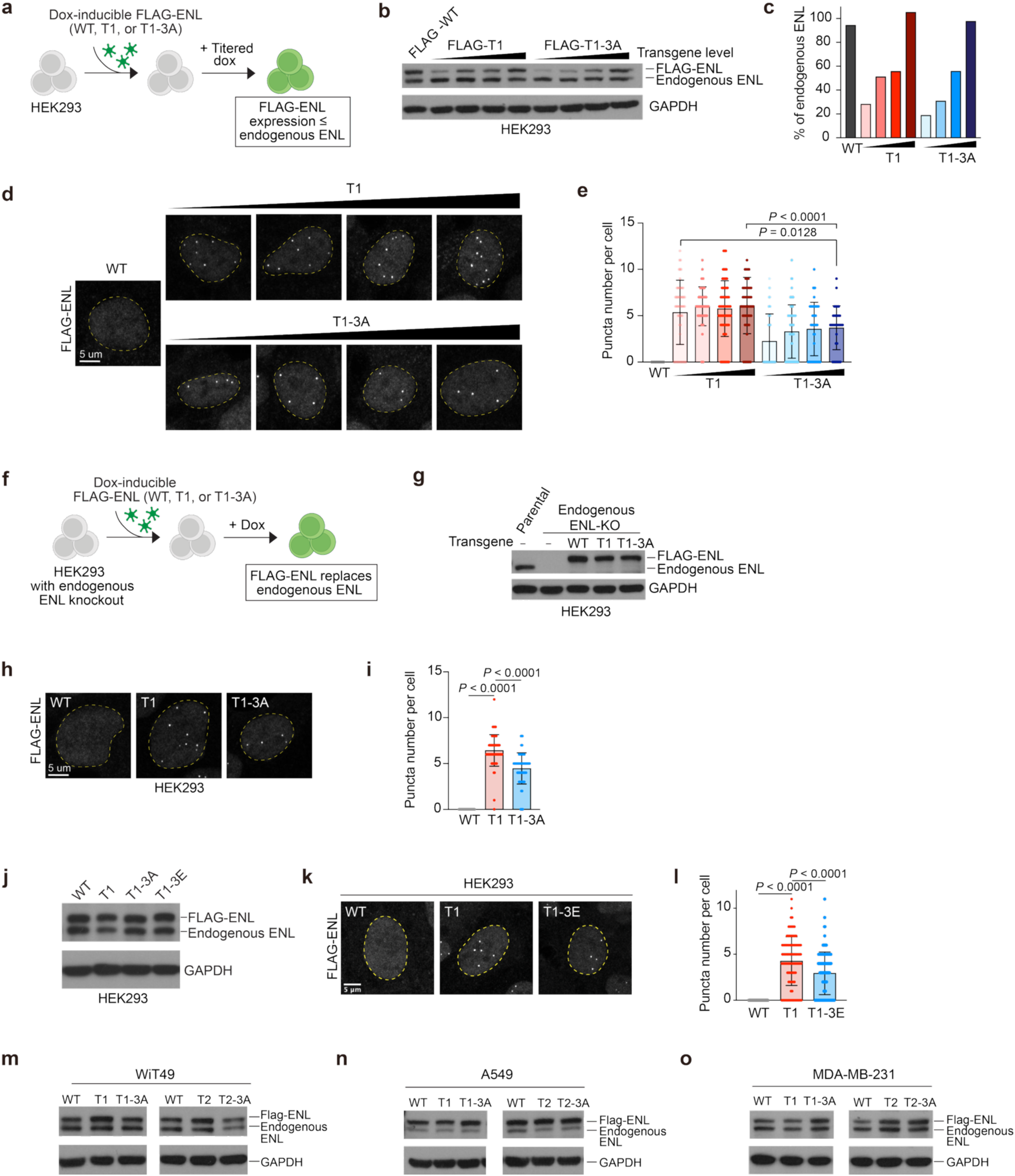
Perturbing RNA interactions impairs ENL mutant condensate formation across cellular contexts. **a**, Schematic of dox-inducible transgene transduction in cells. FLAG-ENL variants are expressed at lower than or close to endogenous levels of ENL. **b**, Western blot showing low to near-endogenous expression of FLAG-ENL variants relative to endogenous ENL expression. WT and T1 data are shown prior in Extended Data Fig. 4b. **c,** Quantification of (**b)**. FLAG-ENL intensity shown as a percentage of endogenous ENL intensity. WT and T1 data are shown prior in Extended Data Fig. 4c. **d**, Representative IF staining of FLAG-ENL variants in HEK293 over increasing T1 expression levels. WT and T1 data are shown prior in Extended Data Fig. 4d. FLAG-WT = ∼endogenous ENL. **e,** Quantification of puncta number per cell. Data are mean ± s.d. *P* values are calculated by two-tailed unpaired Student’s t test. Representative data shown for *n =* 2 replicates. WT and T1 data are shown prior in Extended Data Fig. 4e. FLAG-WT = ∼endogenous ENL. **f,** Schematic of dox-inducible transgene transduction in HEK293 cells with endogenous ENL knockout. FLAG-ENL variants are expressed at near endogenous levels of ENL. **g**, Western blot showing near-endogenous reconstitution of FLAG-ENL variants in ENL-knockout cells. Parental, WT, and T1 data are shown prior in Extended Data Fig. 4h. **h**, Representative IF staining of FLAG-ENL variants in ENL-knockout cells. WT and T1 data are shown prior in Extended Data Fig. 4i. **i,** Quantification of puncta number per cell. Data are mean ± s.d. *P* values are calculated by two-tailed unpaired Student’s t test. Representative data shown for *n =* 2 replicates. WT and T1 data are shown prior in Extended Data Fig. 4j. **j**, Western blot showing near-endogenous expression of FLAG-ENL variants relative to endogenous ENL expression in HEK293. **k,** Representative IF staining of FLAG-ENL variants in HEK293. **l,** Quantification of puncta number per cell. Data are mean ± s.d. *P* values are calculated by two-tailed unpaired Student’s t test. Representative data shown for *n =* 2 replicates. **m-o**, Western blot showing near-endogenous expression of FLAG-ENL variants relative to endogenous ENL expression in WiT49 (**m**), A549 (**n**), and MDA-MB-231 (**o**) cells.

**Extended Data Fig. 6.**
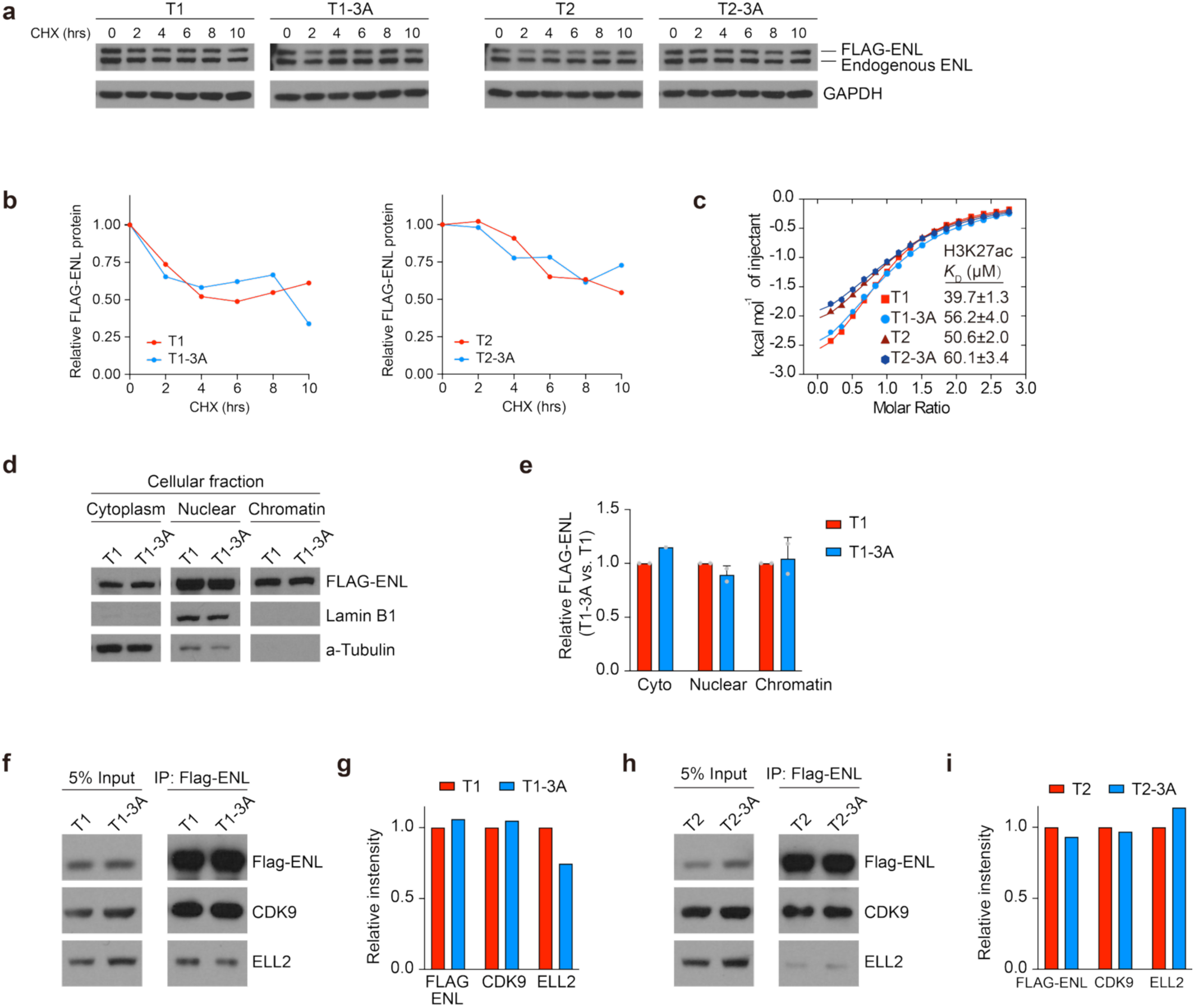
The 3A RNA-binding mutation does not significantly affect other properties of oncogenic ENL mutants. **a**, Western blot showing FLAG-ENL variant expression under cycloheximide (CHX) treatment. **b,** Quantification of (**a**) shows FLAG-ENL variant expression is not altered by the 3A mutation. FLAG-ENL intensity is relative to GAPDH intensity and then normalized to 0 hour treatment timepoint. **c,** Isothermal calorimetry (ITC) using ENL variants and an H3K27ac peptide shows acetyl-binding is not altered by the 3A mutation. *K_D_,* dissociation constant. **d,** Western blot showing cellular fractionation of HEK293 cells expressing FLAG-ENL variants. Lamin B1 and alpha-Tubulin serve as fractionation controls. **e,** Quantification of (**d**) shows FLAG-ENL-T1-3A expression relative to FLAG-T1 expression in each fraction. Data are mean ± s.d. Data from two replicates is shown. **f, h,** Western blot showing immunoprecipitation (IP) of FLAG-ENL variants in HEK293 cells. CDK9 and ELL2 are known ENL interaction partners. **g, i,** Quantification of (**f, h**) shows relative intensity of indicated protein in T1 or T1-3A cells relative to T1 cells.

**Extended Data Fig. 7.**
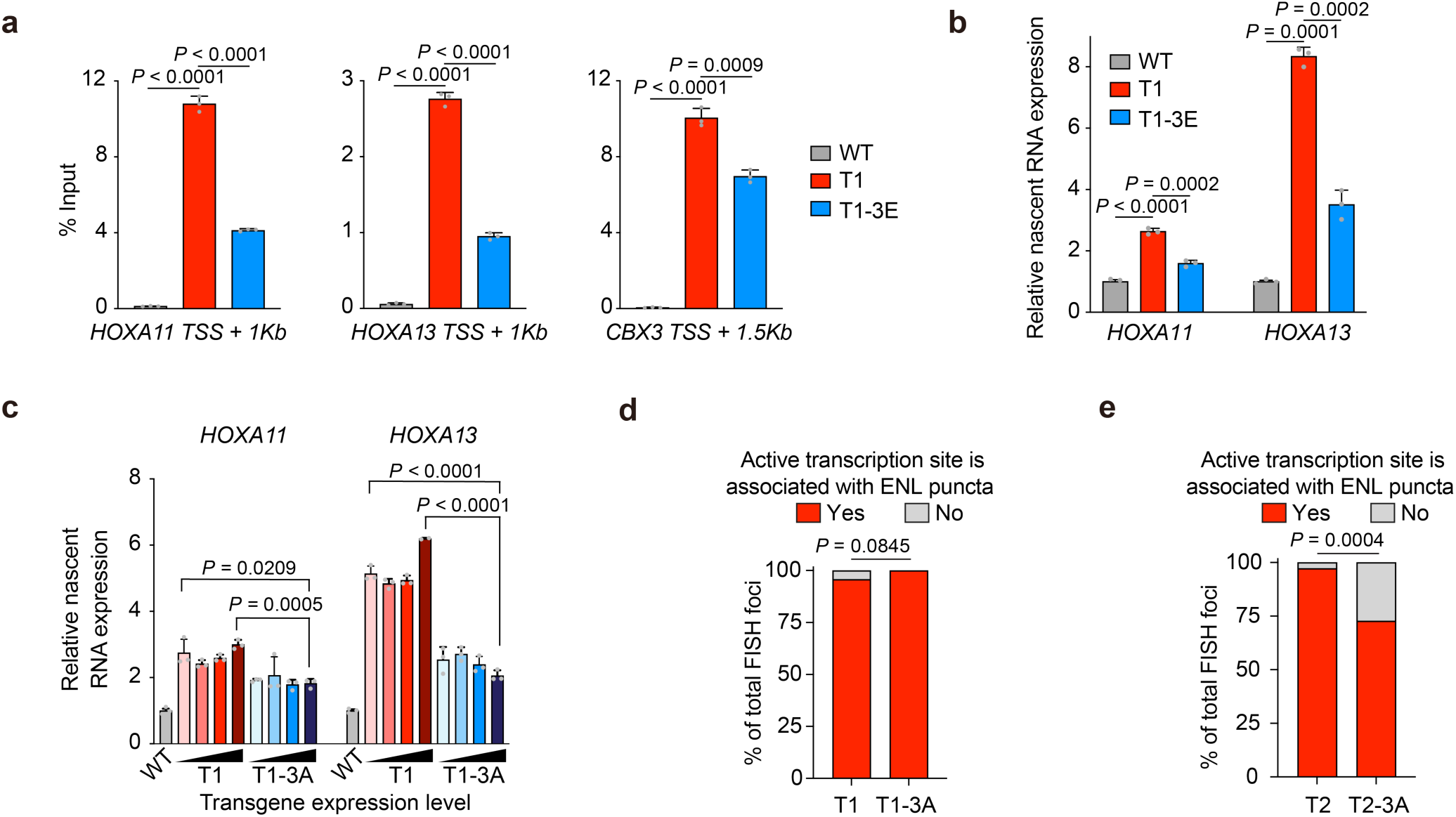
RNA interactions enhance ENL mutant chromatin occupancy at and gene expression of target loci. **a**, ChIP-qPCR of FLAG-ENL variants in HEK293 cells. Data are mean ± s.d. *P* values were calculated by two-tailed unpaired Student’s t test. Triplicate qPCR values are shown. **b,** RT-qPCR showing nascent *HOXA11* or *HOXA13* expression relative to *GAPDH*. Data are mean ± s.d. *P* values were calculated by two-tailed unpaired Student’s t test. Triplicate qPCR values are shown. **c,** RT-qPCR showing nascent *HOXA11* or *HOXA13* expression relative to *18s rRNA*. FLAG-WT = ∼endogenous ENL. Data are mean ± s.d. *P* values were calculated by two-tailed unpaired Student’s t test. Triplicate qPCR values are shown. WT and T1 values are shown prior in Extended Data Fig. 4f. **d, e,** Quantification of percentage of total *HOXA13* RNA FISH foci associated with ENL mutant puncta. *P* values were calculated by two-tailed Fisher’s exact test. Representative data shown for *n =* 2 replicates.

**Extended Data Fig. 8.**
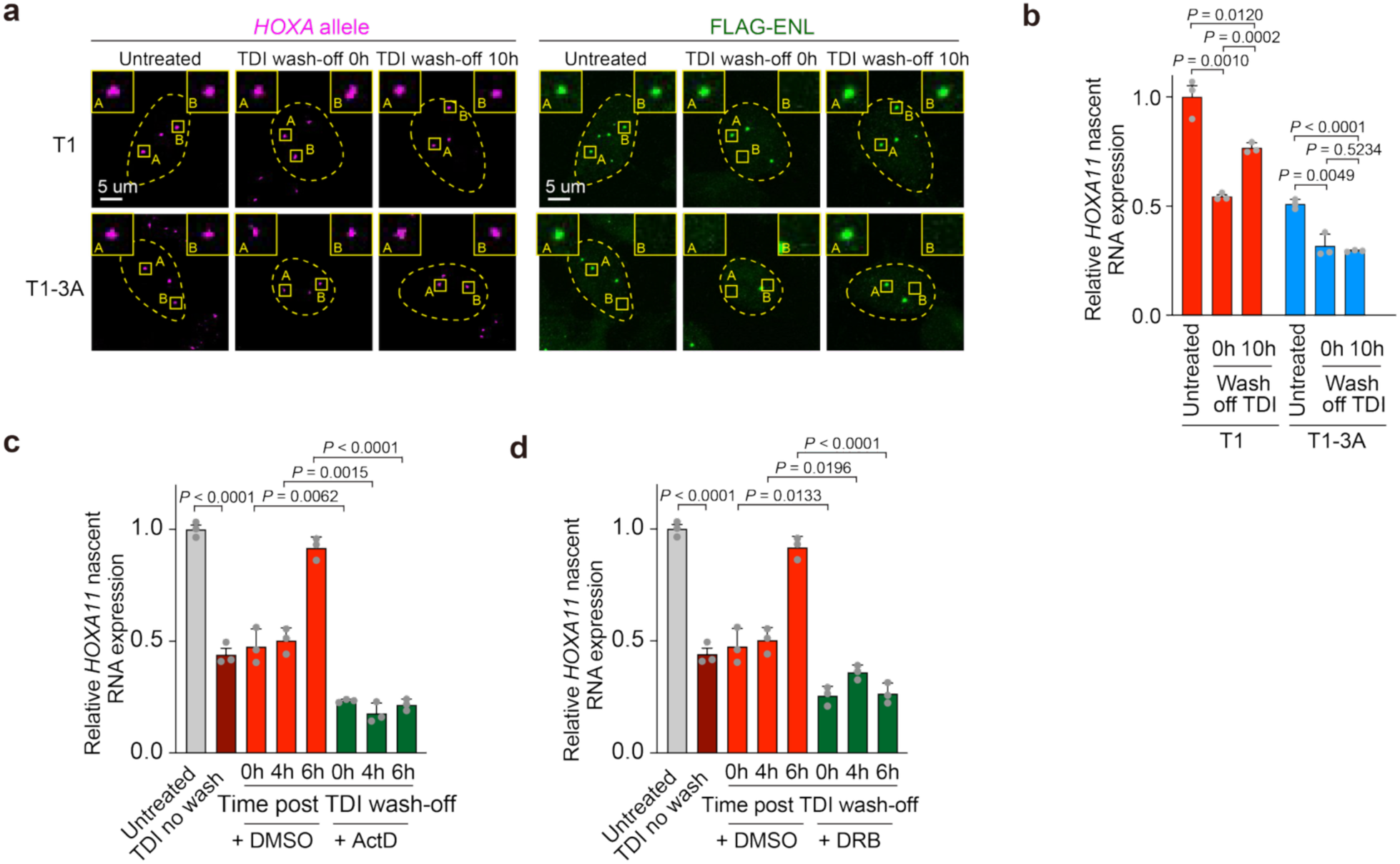
RNA interactions facilitate re-nucleation of ENL mutant condensates at endogenous target loci. **a**, Representative *HOXA* DNA-FISH and FLAG-ENL IF staining in HEK293 cells. Yellow dashed line outlines the nucleus. **b,** RT-qPCR showing nascent *HOXA11* expression relative to *GAPDH*. Data are mean ± s.d. *P* values were calculated by two-tailed unpaired Student’s t test. **c, d,** RT-qPCR showing nascent *HOXA11* expression relative to *18S rRNA*. Data are mean ± s.d. *P* values were calculated by two-tailed unpaired Student’s t test.

**Extended Data Fig. 9.**
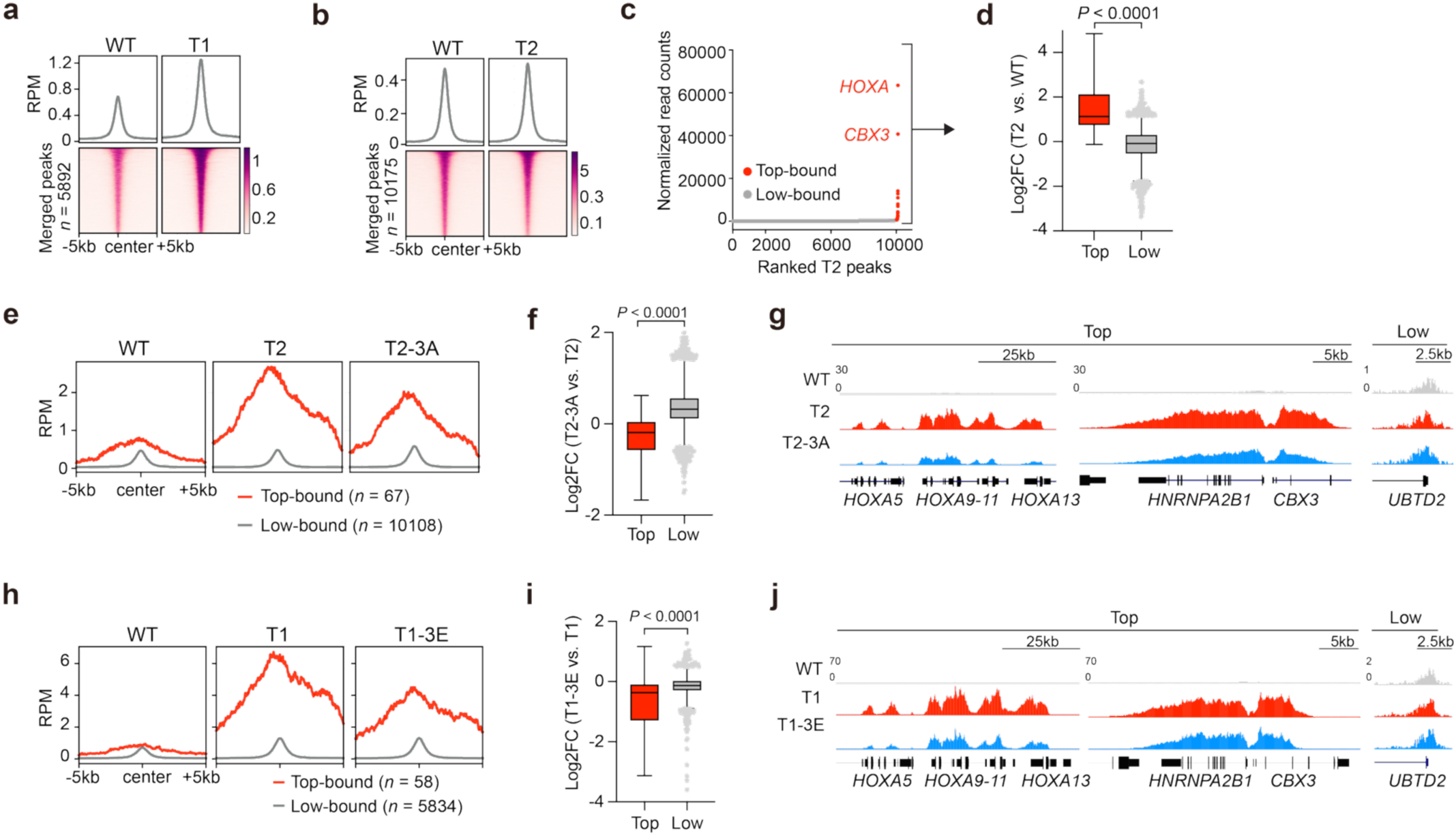
RNA interactions enhance mutant ENL occupancy at condensate-permissive loci in HEK293 cells. **a**, Average (top) and heatmap (bottom) representations of ENL ChIP signal on merged ENL-WT/T1 peaks in HEK293 cells. RPM, reads per million. **b,** Average (top) and heatmap (bottom) representations of ENL ChIP signal on merged ENL-WT/T2 peaks. RPM, reads per million. **c,** ENL-T2 ChIP-seq peaks in HEK293 cells ranked by normalized read count. Top-bound peaks fall to the right of the inflection point and are labeled in red text. Low-bound peaks fall to the left of the inflection point and are labeled in gray. Known condensate-associated targets, *HOXA* and *CBX3*, are labeled in red text. **d,** Box plot showing Log_2_ fold change of T2 vs. WT signal at top- or low-bound sites in HEK293. Top, *n =* 67; Low, *n =* 10108. Center line indicates median and box limits are set to 25^th^ and 75^th^ percentiles. *P* values were calculated by two-tailed unpaired Student’s t test. **e,** Average chromatin occupancy of ENL at top or low-bound sites in HEK293 cells. RPM, reads per million. **f,** Box plot showing Log_2_ fold change of T2-3A vs. T2 signal at top or low-bound sites in HEK293. Top, *n =* 67; Low, *n =* 10108. Center line indicates median and box limits are set to 25^th^ and 75^th^ percentiles. *P* values were calculated by two-tailed unpaired Student’s t test. **g,** Genome browser view of ENL ChIP signal at representative HEK293 top and low-bound genes. **h,** Average chromatin occupancy of ENL at top or low-bound sites in HEK293 cells. RPM, reads per million. **i,** Box plot showing Log_2_ fold change of T1-3E vs. T1 signal at top or low-bound sites in HEK293. Top, *n =* 67; Low, *n =* 10108. Center line indicates median and box limits are set to 25^th^ and 75^th^ percentiles. *P* values were calculated by two-tailed unpaired Student’s t test. **j,** Genome browser view of ENL ChIP signal at representative HEK293 top and low-bound genes.

**Extended Data Fig. 10.**
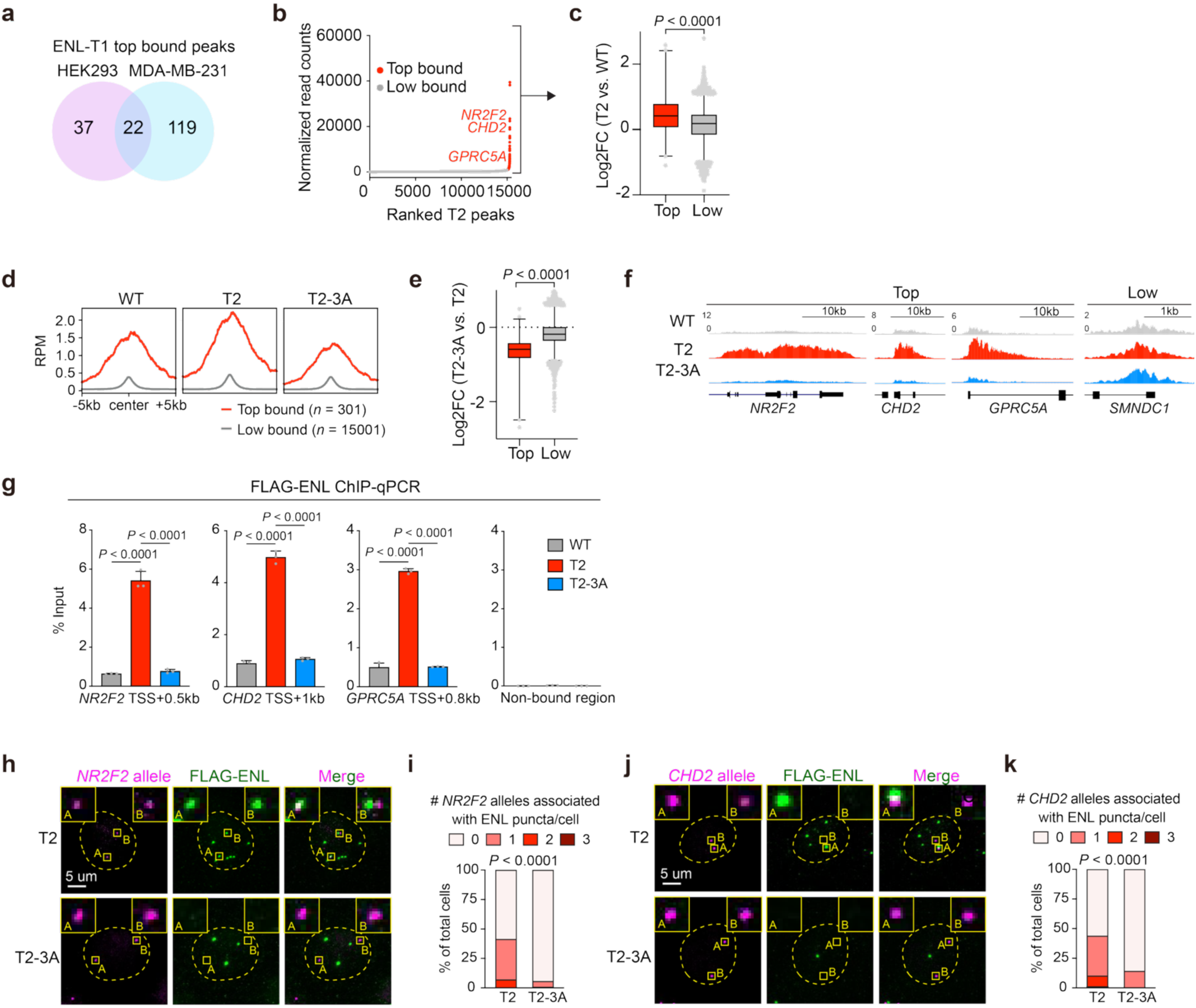
RNA interactions enhance mutant ENL occupancy at condensate-permissive loci in MDA-MB-231 cells. **a**, Venn diagram showing shared or unique top-bound peaks in HEK293 and MDA-MB-231 cells. **b,** ENL-T2 ChIP-seq peaks in MDA-MB-231 cells ranked by normalized read count. Top-bound peaks fall to the right of the inflection point and are labeled in red text. Low-bound peaks fall to the left of the inflection point and are labeled in gray. **c,** Box plot showing Log_2_ fold change of T2/WT signal at top or low-bound sites in MDA-MB-231. Top, *n =* 301; Low, *n =* 15,001. Center line indicates median and box limits are set to 25^th^ and 75^th^ percentiles. *P* values were calculated by two-tailed unpaired Student’s t test. **d**, Average chromatin occupancy of ENL at top or low-bound sites in MDA-MB-231 cells. RPM, reads per million. **e,** Box plot showing Log_2_ fold change of T2-3A vs. T2 signal at top or low-bound sites in MDA-MB-231. Top, *n =* 301; Low, *n =* 15,001. Center line indicates median and box limits are set to 25^th^ and 75^th^ percentiles. *P* values were calculated by two-tailed unpaired Student’s t test. **f,** Genome browser view of ENL ChIP signal at representative top- and low-bound genes in MDA-MB-231 cells. **g,** ChIP-qPCR of FLAG-ENL variants in MDA-MB-231 cells. Data are mean ± s.d. *P* values were calculated by two-tailed unpaired Student’s t test. Triplicate qPCR values are shown. **h, j,** Representative *NR2F2* (**h**) and *CHD2* (**j**) DNA-FISH and FLAG-ENL IF staining in MDA-MB-231 cells. Yellow dashed line outlines the nucleus. Yellow dashed line outlines the nucleus. **i, k,** Quantification of percentage of cells with indicated number of ENL puncta-associated *NR2F2* (**i**) or *CHD2* (**k**) alleles. *P* values were calculated by two-tailed Fisher’s exact test.

**Extended Data Fig. 11.**
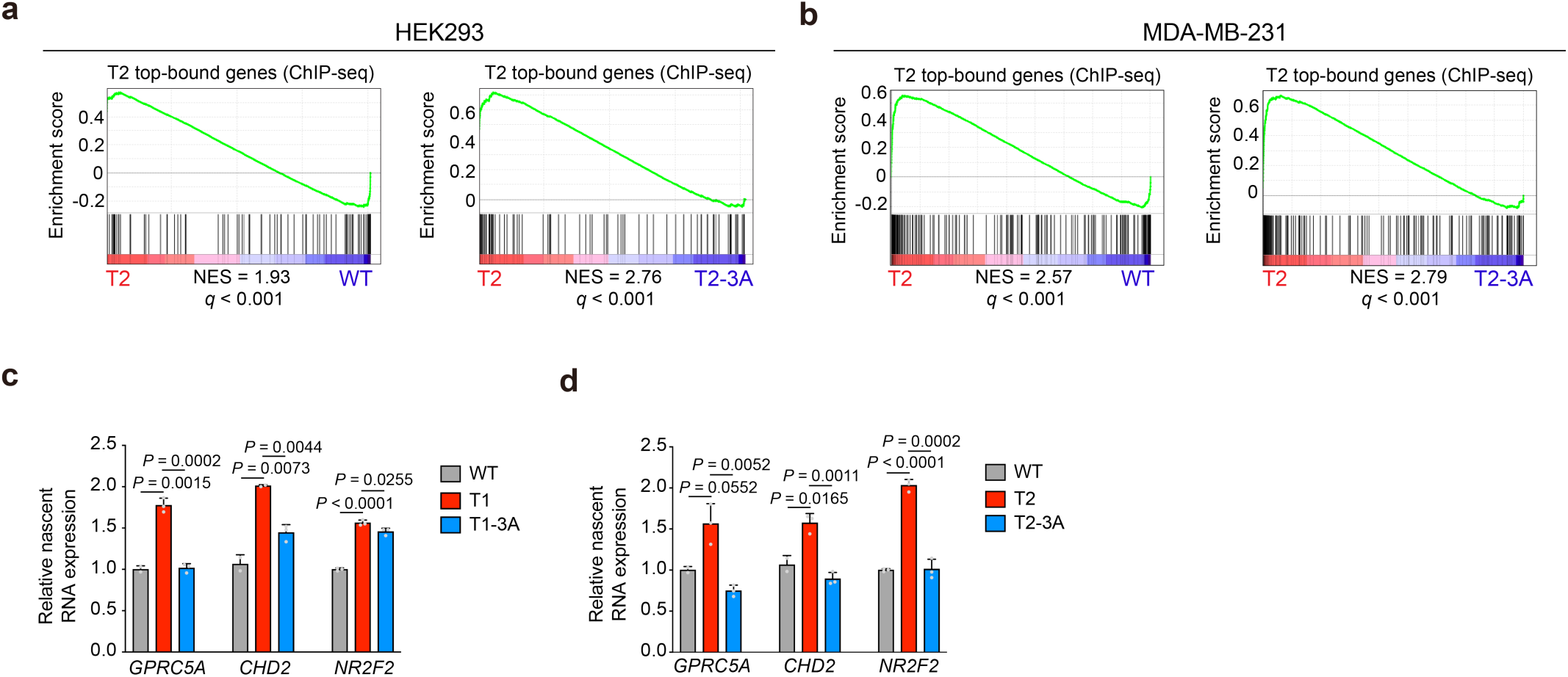
RNA interactions enhance mutant ENL-driven activation of condensate-permissive target loci. **a, b,** GSEA plots showing that top-bound genes (determined by ChIP-seq) are upregulated in T2 compared to WT or to T2-3A in HEK293 (**a**) and MDA-MB-231 (**b**) cells. NES, normalized enrichment score. **c, d,** RT-qPCR showing nascent *GPRC5A*, *NR2F2,* or *CHD2* expression relative to *GAPDH* in MDA-MB-231 cells expressing T1 (**c**) or T2 (**d**) variants. Data are mean ± s.d. *P* values were calculated by two-tailed unpaired Student’s t test. Triplicate qPCR values are shown.

**Extended Data Fig. 12.**
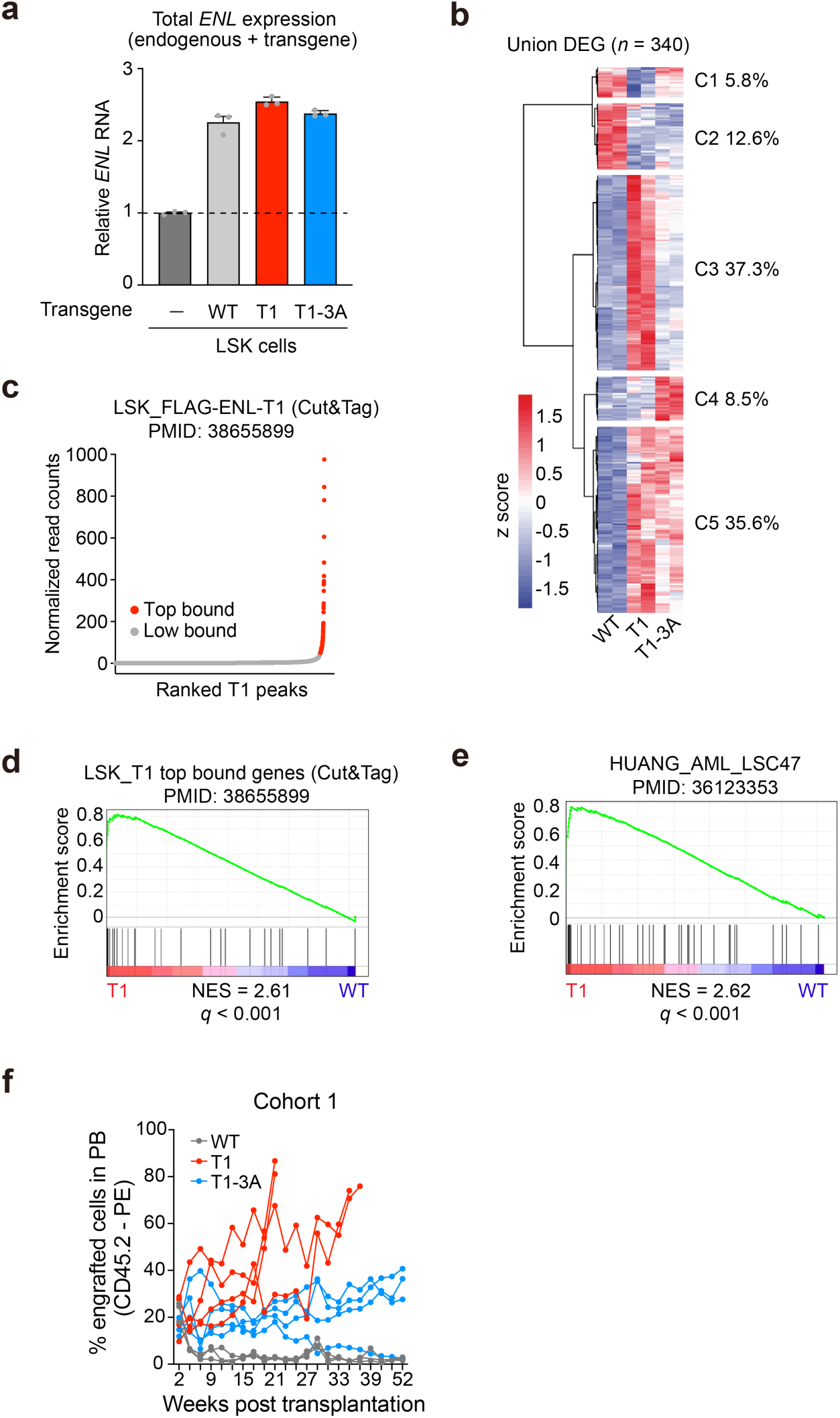
Disrupting RNA binding impairs ENL mutant–driven leukemogenesis. **a**, RT-qPCR showing total human (transgene) and murine (endogenous) *ENL* RNA expression in transduced LSK cells. **b,** Heatmap depicting z-scores of all differentially expressed genes (DEGs). **c,** ENL-T1 Cut&Tag peaks in LSK cells ranked by read count. Top-bound peaks fall to the right of the inflection point and are labeled in red text. Low-bound peaks fall to the left of the inflection point and are labeled in gray. **d,** GSEA plot showing that ENL-T1 top-bound genes are upregulated in T1 vs. WT in LSK cells. NES, normalized enrichment score. **e,** GSEA plot showing that genes associated with AML patients (*n =* 47) are upregulated in T1 vs. WT LSK cells. NES, normalized enrichment score. **g,** Percentage of peripheral blood (PB) composed of cells derived from transplanted LSK. CD45.2, leukocyte marker of donor LSK.

## METHODS

Cell Lines
| Reagent or Resource | Manufacturer/Source | Catalog Number |
| --- | --- | --- |
| HEK293T | ATCC | CRL-3216 |
| HEK293 | ATCC | CRL-1573 |
| WiT49 | Laboratory of Herman Yeger | N/A |
| A549 | Laboratory of Junwei Shi | N/A |
| MDA-MB-231 | Laboratory of Yibin Kang | N/A |

Antibodies
| Reagent or Resource | Manufacturer | Catalog Number |
| --- | --- | --- |
| Brilliant Violet 421™ anti-mouse CD3 Antibody | Biolegend | 100227 |
| Brilliant Violet 421™ anti-mouse CD19 Antibody | Biolegend | 115549 |
| Brilliant Violet 421™ anti-mouse/human CD11b Antibody | Biolegend | 101251 |
| Brilliant Violet 421™ anti-mouse/human CD45R/B220 Antibody | Biolegend | 103240 |
| Brilliant Violet 421™ anti-mouse TER-119/Erythroid Cells Antibody | Biolegend | 116233 |
| Brilliant Violet 421™ anti-mouse Ly-6G/Ly-6C (Gr-1) Antibody | Biolegend | 108433 |
| Brilliant Violet 421™ anti-mouse NK-1.1 Antibody | Biolegend | 108741 |
| Brilliant Violet 421™ anti-mouse CD11c Antibody | Biolegend | 117343 |
| Brilliant Violet 421™ anti-mouse TCR $\gamma/\delta$ Antibody | Biolegend | 118119 |
| Brilliant Violet 421™ anti-mouse TCR $\beta$ chain Antibody | Biolegend | 109229 |
| PerCP/Cyanine5.5 anti-mouse Ly-6A/E (Sca-1) | Biolegend | 108124 |
| PE anti-mouse CD117 (c-Kit) | Biolegend | 105808 |
| CD45.1 mouse anti-mouse APC/Cy7 | Fisher Scientific | BDB560579 |
| CD45.2 mouse anti-mouse PE | Fisher Scientific | BDB560695 |
| Flag | Sigma | F1804 |
| MLLT1/ENL | Millipore Sigma | ABE2596 |
| GAPDH | Cell Signaling Technology | 2118 |
| CDK9 | Cell Signaling Technology | 2316S |
| ELL2 | Bethyl | A302-505A-1 |
| LaminB1 | Abcam | ab16048 |
| α-Tubulin | Santa Cruz | sc-5286 |
| Anti-mouse IgG, HRP-linked Antibody | Cell Signaling Technology | 7074S |
| Anti-rabbit IgG, HRP-linked Antibody | Cell Signaling Technology | 7076S |
| Goat anti-Rabbit IgG (H+L) Cross-Adsorbed Secondary Antibody, Alexa Fluor 488 | Invitrogen | A32731 |
| Goat anti-Mouse IgG (H+L) Cross-Adsorbed Secondary Antibody, Alexa Fluor 488 | Invitrogen | A32723 |
| Goat anti-Rabbit IgG (H+L) Cross-Adsorbed Secondary Antibody, Alexa Fluor 568 | Invitrogen | A-11036 |
| Goat anti-Mouse IgG (H+L) Cross-Adsorbed Secondary Antibody, Alexa Fluor 568 | Invitrogen | A11031 |
| Goat anti-Rabbit IgG (H+L) Cross-Adsorbed Secondary Antibody, Alexa Fluor 647 | Invitrogen | A-21245 |
| Goat anti-Mouse IgG (H+L) Cross-Adsorbed Secondary Antibody, Alexa Fluor 647 | Invitrogen | A-21236 |

Commercial Assays
| Reagent or Resource | Manufacturer | Catalog Number |
| --- | --- | --- |
| HisTrap HP His tag protein pur columns | Cytiva | 29051021 |
| Superdex 75 10/300 GL columns | Cytiva | 28920504 |
| Pierce microcentrifuge spin columns | Thermo Fisher | 89879 |
| RNA Clean & Concentrator Kit | Zymo Research | R1013 |
| QIAquick PCR Purification Kit | Qiagen | 28104 |
| Vazyme Universal DNA Library Prep Kit for Illumina V4 | Vazyme | ND610-04 |
| Qubit 1X dsDNA High Sensitivity Kit | Invitrogen | Q33231 |
| Agilent D1000 Screen Tape System | Agilent | N/A |
| RNeasy Mini Kit | Qiagen | 74104 |
| High-Capacity cDNA Reverse Transcription Kit | Applied Biosystems | 4368814 |
| NEBNext Poly(A) mRNA Magnetic Isolation Module | New England Biolabs | E7490L |
| NEBNext Ultra™ II RNA Library Prep Kit for Illumina | New England Biolabs | E7770L |
| NEBNext Multiplex Oligos for Illumina | New England Biolabs | E7335S |
| EasySep Mouse CD117 (cKIT) Positive Selection Kit | Stem Cell Technologies | 18757 |

Media, chemicals, and other reagents
| Reagent or Resource | Manufacturer | Catalog Number |
| --- | --- | --- |
| DMEM with L-Glutamine, 4.5g/L Glucose and Sodium Pyruvate | Corning | MT10013CV |
| Eagle's Minimum Essential Medium (EMEM) | Corning | MT10009CV |
| DMEM/F-12 | Sigma | D6421 |
| IMDM + GlutaMAX | Gibco | 31980030 |
| Opti-MEM | Gibco | 31985070 |
| Hank's Balanced Salt Solution (HBSS) | Gibco | 14-170-112 |
| Fetal Bovine Serum | Cytiva | SH103IH2540 |
| Penicillin/Streptomycin | Gibco | 15070063 |
| Cycloheximide | Cell Signaling Technology | 2112S |
| MethoCult | Stem Cell Technologies | 03234 |
| Recombinant murine Flt3 ligand | PeproTech | 250-31L |
| Recombinant murine SCF | PeproTech | 250-03 |
| Recombinant murine TPO | PeproTech | 315-14 |
| Recombinant murine IL6 | PeproTech | 216-16 |
| Recombinant murine IL3 | PeptoTech | 200-03 |
| RBC Lysis Buffer (10x) | BioLegend | 420301 |
| Zombie Aqua Fixable Viability Kit | BioLegend | 423101 |
| SYTOX Blue Dead Cell Stain | Invitrogen | S34857 |
| 2-Mercaptoethanol | BioRad | 1610710 |
| L-glutamine | Gibco | A2916801 |
| PEI MAX | Polysciences | 24765 |
| Polyethylene Glycol 8000 (PEG) | Fisher Scientific | BP233-100 |
| Polybrene | Millipore Sigma | TR-1003-G |
| Doxycycline | Sigma | D9891-1G |
| RetroNectin Recombinant Human Fibronectin Fragment | Takara Bio | T100B |
| Bovine Serum Albumin | Sigma Aldrich | A7906-10G |
| 16% Paraformaldehyde Aqueous | Electron Microscopy Sciences | 15710 |
| Tween-20 | Fisher Scientific | BP337500 |
| Triton-X-100 | Fisher Scientific | BP151500 |
| Goat serum | Cell Signalling Technology | 5425s |
| Mounting Medium with DAPI - Aqueous, Fluoroshield | Abcam | ab104139 |
| PowerUp™ SYBR™ Green Master Mix for qPCR | Applied Biosystems | A25778 |
| Glycine | Fisher Scientific | BP3815 |
| Tris Base | Fisher Scientific | BP152-1 |
| Blotting grade blocker | BioRad | 1706404 |
| SDS | Thermo Fisher | BP8200500 |
| Pierce ECL Western Blotting Substrate | Thermo Scientific | 32106 |
| Tris-HCl | Fisher Scientific | BP153-500 |
| EDTA | Sigma Aldrich | E5134-500G |
| NaDOC | Sigma Aldrich | 35915 |
| NaCl | Fisher Scientific | BP358-1 |
| Sarkosyl | Sigma Aldrich | 61739 |
| DTT | Sigma Aldrich | D9163-5G |
| cOmplete EDTA-free Protease Inhibitor Cocktail | Sigma-Aldrich | 11836170001 |
| Dynabeads Protein A for IP | Invitrogen | 10001D |
| LiCl | Sigma-Aldrich | 213233-500G |
| NP-40 | BioVision | 2111-100 |
| RNase A | Millipore Sigma | 9001-99-4 |
| Proteinase K | Thermo Scientific | EO0491 |
| Poly-L-lysine solution | Sigma Aldrich | P8920 |
| 20X SSC buffer | Corning | 46-020-CM |
| Formamide | Fisher Scientific | BP227-500 |
| Dextran Sulfate | Sigma Aldrich | D8906 |
| PVSA solution | Sigma Aldrich | 278424 |
| Hoechst 33342 solution, 20 mM | Thermo Scientific | 62249 |
| SlowFade Gold Antifade Mountant | Invitrogen | S36936 |
| Glycerol | Fisher Scientific | P2291 |
| RNase inhibitor (Murine) | Beyotime | R0102 |
| 10X TBE RNase-free | MREDA | M212499 |
| APS | Sigma Aldrich | 7727-54-0 |
| TEMED | Sigma Aldrich | 110-18-9 |
| 30%Arc-Bis RNase-free | Macklin | A917692 |
| Imidazole | Sigma Aldrich | 12399 |
| D-Desthiobiotin | IBA Life Sciences | 2-1000-002 |
| Isopropyl β-D-1-thiogalactopyranoside | Sigma Aldrich | 10724815001 |
| Kanamycin | Fisher Scientific | BP906-5 |
| Ni IDA Beads 6FF | Smart-Lifesciences | SA00501L |
| Strep-4FF beads | Smart-Lifesciences | SA053100 |
| Alexa 488 Fluor™ 488 5-TFP | Invitrogen | A30005 |
| HEPES-Na | Sigma-Aldrich | H7006 |
| H3<sub>34-28</sub>K27ac | This paper | SciLight<br>Biotechnology, LLC |
| 5' CY5 HOXA11 ssRNA 15nt/30nt | This paper | hippobio |
| 5' CY5 HOXA11 ssDNA 15nt/30nt | This paper | xianghongbio |
| DEPC | Sigma-Aldrich | 1609-47-8 |
| BL21 (DE3) competent cells | Millipore Sigma | 69450-3 |
| HaloLink resin | Promega | G1913 |
| Bovine serum albumin (BSA) fraction V | Sigma-Aldrich | 126609-5GM |
| 50X protease inhibitor cocktail | Promega | G6521 |
| TURBO DNase | Invitrogen | AM2238 |
| RNase Inhibitor, murine | New England Biolabs | M0314S |
| Ambion RNase I | Invitrogen | AM2294 |
| FastAP Thermosensitive Alkaline Phosphatase | Thermo Scientific | EF0651 |
| T4 Polynucleotide Kinase (PNK) | New England Biolabs | M0201S |
| T4 RNA ligase high concentration, T4 RNA ligase reaction buffer, 50% PEG8000, ATP | New England Biolabs | M0204 |
| Dynabeads MyOne Silane | Invitrogen | 37002D |
| Maxima H Minus Reverse Transcriptase | Thermo Scientific | EP0752 |
| Exol | New England Biolabs | M0293S |
| Instant Sticky-end Ligase Master Mix | New England Biolabs | M0370S |
| AMPure XP beads | Beckman Coulter | A63881 |
| Janelia Fluor 657 (JF657) | Laboratory of Luke Lavis | N/A |
| Ribonucleoside vanadyl complexes (RVC) | Sigma Aldrich | R3380-5ML |
| Actinomycin D | Millipore Sigma | A9415-2MG |
| 5,6-Dichlorobenzimidazole 1-β-D-ribofuranoside (DRB) | Millipore Sigma | D1916 |
| Triptolide | Sigma | T3652-1MG |
| Alpha-Amanitin | Sigma | A2263-1MG |

Recombinant DNA
| Reagent or Resource | Source | Catalog Number |
| --- | --- | --- |
| VSVG | Addgene | 8454 |
| PAX2 | Addgene | 12260 |
| pCL-ECO | Addgene | 12371 |
| pLenti-tetON-miniCMV-3xFLAG-2xHA-hsENL-WT-NLS | Song *et al.*, *Mol Cell*, 2022 | N/A |
| pLenti-tetON-miniCMV-3xFLAG-2xHA-hsENL-T1-NLS | Song *et al.*, *Mol Cell*, 2022 | N/A |
| pLenti-tetON-miniCMV-3xFLAG-2xHA-hsENL-T1(K61A/R63A/K67A)-NLS ('T1-3A') | This paper | N/A |
| pLenti-tetON-miniCMV-3xFLAG-2xHA-hsENL-T1(K61E/R63E/K67E)-NLS ('T1-3E') | This paper | N/A |
| pLenti-tetON-miniCMV-3xFLAG-2xHA-hsENL-T2-NLS | Song *et al.*, *Mol Cell*, 2022 | N/A |
| pLenti-tetON-miniCMV-3xFLAG-2xHA-hsENL-T2(K61A/R63A/K67A)-NLS ('T2-3A') | This paper | N/A |
| pLenti-tetON-miniCMV-Halo-3xFLAG-hsENL-WT-NLS | Song *et al.*, *Mol Cell*, 2022 | N/A |
| pLenti-tetON-miniCMV-Halo-3xFLAG-hsENL-T1-NLS | Song *et al.*, *Mol Cell*, 2022 | N/A |
| pLenti-tetON-miniCMV-Halo-3xFLAG-hsENL-T1(K61A/R63A/K67A)-NLS ('T1-3A') | This paper | N/A |
| pLenti-tetON-miniCMV-Halo-3xFLAG-hsENL-T2-NLS | Song *et al.*, *Mol Cell*, 2022 | N/A |
| pLenti-tetON-miniCMV-Halo-3xFLAG-hsENL-T2(K61A/R63A/K67A)-NLS ('T2-3A') | This paper | N/A |
| MSCV-3xFLAG-HA-hsENL-WT-PGK-puro-IRES-GFP | Liu *et al.*, *Cancer Discov*, 2024 | N/A |
| MSCV-3xFLAG-HA-hsENL-T1-PGK-puro-IRES-GFP | Liu *et al.*, *Cancer Discov*, 2024 | N/A |
| MSCV-3xFLAG-HA-hsENL-T1(K61A/R63A/K67A)-PGK-puro-IRES-GFP | This paper | N/A |
| pET28a-2*strep-ENL-WT-6*his | Song *et al.*, *Mol Cell*, 2022 | N/A |
| pET28a-2*strep-ENL-T1-6*his | Song *et al.*, *Mol Cell*, 2022 | N/A |
| pET28a-2*strep-ENL-T1(K61A/R63A/K67A)-6*his | This paper | N/A |
| pET28a-2*strep-ENL-T1-IDR(K/Q<sub>21</sub>)-6*his | Song *et al.*, *Mol Cell*, 2022 | N/A |
| pET28a-6*his-ENL-WT(1-148) | Song *et al.*, *Mol Cell*, 2022 | N/A |
| pET28a-6*his-ENL-T1(1-151) | Song *et al.*, *Mol Cell*, 2022 | N/A |
| pET28a-6*his-ENL-T1(K61A/R63A/K67A)(1-151) | This paper | N/A |

Software and algorithms
| Reagent or Resource | Manufacturer | Catalog Number |
| --- | --- | --- |
| FIJI | Schindelin *et al.*, 2012 | imagej.net/software/fiji/ |
| GraphPad Prism | GraphPad | graphpad.com |
| Imaris 9.3.1 | Oxford Instruments | imaris.oxinst.com |
| Origin 7.0 | OriginLab | originlab.com |
| PyMOL 3.1 | PyMOL | pymol.org |
| AlphaFold3 Server | N/A | PMID: 38718835 |
| FlowJo | BD Biosciences | flowjo.com |
| GSEA_4.3.2 | N/A | PMID: 16199517;<br>PMID: 12808457 |

### Protein purification

For full-length ENL protein purification, N-terminal 2 X streptavidin tags and C-terminal 6 X histidine tags were added to the pET28b expression vector containing the ENL-WT CDS (residues 1-559; NP_005925.2). The T1 mutation (p. 117_118insNHL) and RNA-binding residue mutations (K61A, R63A, and K67A) were subsequently added. Proteins were expressed in BL21(DE3) *E. coli* cells induced with 0.4mM IPTG (Isopropyl β-D-thiogalactopyranoside) for 16 h at 16°C. Cells were harvested and resuspended in lysis buffer (25 mM Tris-HCl pH 7.5, 500 mM NaCl, 5% glycerol, 50 mM imidazole, 1 mM PMSF). After lysis using an Emulsiflex C3 (Avestin) high-pressure homogenizer, crude lysate was clarified by centrifugation. Clarified lysate was loaded onto a HisTrap column and washed with wash buffer (30 mM Tris-HCl pH 7.5, 1 M NaCl, 5% glycerol). Protein was eluted using lysis buffer supplemented with 500 mM Imidazole, concentrated, and buffer-exchanged to strep buffer (25 mM HEPES-Na pH 7.4, 500 mM NaCl, 5% glycerol, 1 mM EDTA, 2 mM β-mercaptoethanol) to decrease the imidazole concentration. After buffer exchange, the protein was incubated with strep-4FF beads, washed with strep buffer, and eluted using strep buffer supplemented with 5 mM desthiobiotin.

For purification of the ENL YEATS (residues 1-148) domain, N-terminal 6 X histidine tags were added to the pET28b vector containing the ENL-WT YEATS domain CDS (residues 1-148). The T1 mutation (p. 117_118insNHL) and RNA-binding residue mutations (K61A, R63A, and K67A) were subsequently added. Proteins were expressed in BL21(DE3) *E. coli* cells induced with 0.4mM IPTG for 16h at 16°C. Cells were harvested and resuspended in lysis buffer. After lysis using an Emulsiflex C3 high-pressure homogenizer, crude lysate was clarified by centrifugation. Clarified lysate was loaded onto a HisTrap column and washed with wash buffer (30 mM Tris-HCl pH 7.5, 1 M NaCl, 5% glycerol). Protein was eluted with a linear imidazole gradient from 20 mM to 500 mM. Protein was further purified and polished over a Superdex 75 10/300 gel filtration column using AKTA Purifier 10 systems (GE Healthcare). The monomer peak in elution buffer (500 mM NaCl, 25 mM Tris-HCl pH 7.5, and 2 mM β-mercaptoethanol) was pooled, concentrated to about 10 mg/mL, and stored at −80°C.

### AlphaFold predictions and analysis

We used the Alphafold-Multimer (v3) to predict the structure of the ENL YEATS-RNA complex. The human ENL 1-142 T1 sequence was submitted as the protein sequence, and the first 15 nucleotides of 104 randomly selected transcripts were chosen as input RNA sequences (Supplementary Table 1). Each model was predicted using one ENL YEATS domain and one RNA fragment. For all predicted results, the top-scoring relaxed model was selected, with an average iPTM score > 0.6 being inferred as putative hits. Diagnostic plots (PAE plot, PLDDT plot, and sequence coverage) and generated structures were manually checked. The regions characterizing the interaction between ENL and RNA were extracted from the PAE matrix, and the minimum PAE corresponding to each ENL residue was calculated and normalized. The PAE values corresponding to each residue were summed and mapped onto the structure for visualization.

To examine the impact of the 3A mutation on ENL:RNA interactions, we performed the same Alphafold modelling with the T1-3A YEATS domain sequence and the 104 RNA sequences. For the 104 T1:RNA models and 104 T1-3A:RNA models, we measured the shortest possible distance between each of ENL’s putative RNA binding residues (K61, R63, and K67) and the RNA molecule. We then plotted the percentage of models that show each individual residue as being within 3, 4, or 5 Å of the RNA molecule.

### Electrophoretic mobility shift assay (EMSA)

The nucleic acids used in the experiment were all 5’ cy5 labeled. ssRNA was first incubated at 95°C for 5 minutes and then quickly cooled on ice for 5 minutes. RNA refolding buffer (10 mM Bis-Tris pH 6.7, 50 mM KCl, 10 mM MgCl_2_; 1X final) was then added to the RNA on ice. To refold, the RNA was transferred to a cold metal block and folded by warming to room temperature (∼25°C) for 30 minutes. For dsDNA, the cy5 labeled DNA oligo was mixed with an equal amount of unlabeled complementary oligo in DNA annealing buffer (10 mM Tris-HCl pH 8.0, 50 mM NaCl, 1 mM EDTA,1 X final), heated at 90°C for 3 minutes, and then cooled slowly to room temperature. The prepared nucleic acids were incubated with purified ENL protein in RNase-free ultra-low adhesion PCR tubes. The reactions contained 1 X binding buffer (1x PBS with 5% glycerol, 2 mM β-ME, and 0.4 U/μL Murine RNase inhibitor (Beyotime)). The reactions were loaded on a 6% native polyacrylamide gel (29:1 Acrylamide: Bis-acrylamide) and electrophoresed at 100 V for 1 hour in 0.5x TBE. The gel was imaged and photographed using a Tanon gel imaging system with a 605 nm light source. The experiment was performed with at least three technical replicates and three biological replicates. All reagents and equipment used in the experiment were RNase-free.

Upper (bound) and lower (unbound) band intensities were quantified in ImageJ. To obtain the fraction of bound nucleic acids in each lane, the intensity of the most upper band(s) was divided by the intensity of the lower band in the free nucleic acid lane as a representation of the total amount of nucleic acid per lane. For 15 nt experiments, the upper band(s) were defined by the top most band. For 70 nt RNA experiments, the upper band(s) were defined by the region encompassing all bands above the free RNA band. Per plot, fractions from three technical replicates are shown. 3-6 biological replicates were performed for each experiment. Shown is one representative biological replicate per experiment.

### *In vitro* droplet formation assay

Droplet formation assay was performed in the droplet formation buffer (1X PBS with 5% glycerol and 2 mM β-ME). Proteins were fluorescently labeled for microscopy using Microscale Protein Labeling kits from Thermo Scientific: Alexa Fluor 488 (A30005). Synthetic nucleic acids labeled with 5’ cy5 were used in the experiment. Labeled proteins were added to unlabeled proteins at a 1:10 molar ratio. Droplets were assembled in 384 low-binding multi-well 0.17-mm microscopy plates (384-well microscopy plates) (In Vitro Scientific) and sealed with optically clear adhesive film. For the groups with RNase treatment, RNase A powder is pre-dissolved in droplet formation buffer. After ENL and RNA have been incubated in droplet formation buffer, RNase A is added to achieve a final concentration of 1 µg/mL and incubated at room temperature for 5 minutes. All reagents used in the experiment were RNase-free. After quick centrifugation, droplets were imaged under Olympus FV1200 using a 100 X oil objective. The results were processed by Imaris 9.3.1 (Oxford Instrument). Total droplet area was defined by and plotted as the total area of a field occupied by droplets. Total RNA signal per droplet is defined by the sum of RNA channel pixel intensities within a mask drawn over each individual droplet.

### Isothermal titration calorimetry (ITC)

All ITC titrations were performed at 25°C using MicroCal iTC200 system (Malvern). Both synthetic histone H3 peptides (H3_24-28_K27ac) and the recombinant ENL YEATS domain proteins were extensively dialyzed against ITC buffer (25 mM Tris-HCl pH 7.5, 500 mM NaCl, 5 % glycerol and 2 mM β-ME). Before titration, all samples were centrifuged at 13,000 rpm at 4°C for ∼10-15 minutes to remove potential aggregates. ITC cells were rinsed with buffer for several times before protein solution was added to the cell. Cell contents were pipetted several times to mix with any trace buffer remaining in the cell. A small volume of protein solution was removed from the cell for concentration re-check using a NanoDrop. Each ITC titration consisted of 17 successive injections with 0.5 μL for the first and 2.4 μL for the rest. The intervals between injections were 150 seconds, and the stirring speed was 750 rpm. Histone H3 peptides (2.0-2.2 mM) were titrated into ENL YEATS domain proteins (0.15 mM) in all experiments. The resultant ITC curves were processed using the Origin (v.7.0) software (OriginLab) in accordance with the “One Set of Sites” fitting model. First injection data was always excluded from the analysis.

### Human cell lines

All cell lines used in this study were cultured in a humidified incubator maintained at 37°C and 5% CO_2_ and regularly tested mycoplasma negative. All cell lines not purchased from ATCC were subjected to STR authentication testing prior to any experimentation. HEK293T, A549, MDA-MB-231 were cultured in DMEM supplemented with 10% FBS and 100 U/mL penicillin/streptomycin. HEK293 cells were cultured in EMEM supplemented with 10% FBS and 100 U/mL penicillin/streptomycin. WiT49 cells were cultured in DMEM/F-12 supplemented with 10% FBS and 100 U/mL penicillin/streptomycin.

### Lentivirus and retrovirus generation

HEK293T cells were used for all virus packaging. For lentivirus generation, cells were seeded 1 day prior to transfection. For transfection, lentiviral packaging plasmids VSVG and PAX2 were combined with the plasmid of interest and Polyethylenimine MAX (PEI MAX) in Opti-MEM at a ratio of 1:4, total DNA (ug): PEI (uL). These components were incubated to allow for complex formation and added to the HEK293T cells. ∼6h after adding the mix, transfection media was removed and replaced with fresh culture media. The next day, virus was collected and fresh culture media was added back to the cells. A total of 5 virus collections occurred over the course of 72h post-transfection. For retrovirus generation, cells were seeded 1 day prior to transfection. For transfection, retrovirual packaging plasmid pCL-ECO was combined with the plasmid of interest and Polyethylenimine MAX (PEI MAX) in Opti-MEM at a ratio of 1:3, total DNA (ug): PEI (uL). These components were incubated to allow for complex formation and added to the HEK293T cells. ∼6h after adding the mix, transfection media was removed and replaced with fresh culture media. The next day, virus was collected and fresh culture media was added back to the cells. A total of 5 virus collections occurred over the course of 72h post-transfection. When necessary, virus was concentrated 100x with 5x PEG8000 solution.

### Human cell line viral transduction and transgene expression

HEK293 and WiT49 stable cell lines were generated by incubating cells (∼70% confluent) with concentrated lentivirus and 10 ug/mL polybrene overnight (∼16h). A549 and MDA-231 stable cell lines were generated by adding concentrated lentivirus and 10 ug/mL polybrene to cells (∼70% confluent), performing spinfection (2500 rpm for 1h), and incubating overnight. When applicable, selection regents were added 24-48 hours post-infection. Cells containing doxycycline-inducible transgenes (i.e., ENL) were incubated for 48h with a pre-determined concentration of doxycycline such that exogenous ENL expression was similar to that of endogenous ENL as measured by protein levels in Western blot.

### Generation of T1 knock-in HEK293 cell line

Insertion of the T1 mutation was conducted via electroporation of CRISPR/Cas9 and a homologous recombination construct in exon 4 of *ENL* in HEK293 cells previously containing a Halo-tag knockin 5’ to the *ENL* CDS. Single clones were isolated. To verify efficient T1 mutation knock-in, genomic DNA was isolated from cells and a 200bp region surrounding the knock-in target location was amplified by PCR. The PCR product was gel purified and subjected to Sanger sequencing and full CRISPR sequencing to analyze knock-in frequency. CRISPR sequencing was performed by the Massachusetts General Hospital DNA core. The clone shown is homozygous for the T1 mutation. Primers used for PCR amplification and Sanger sequencing are listed in Supplementary Table 11.

### CLAP pull-down and RNA isolation

CLAP-sequencing was conducted according to published protocols from the Dr. Mitchell Guttman lab^35^ with some modifications. For CLAP experiments, HEK293 cells stably expressing doxycycline-inducible Halo-tagged ENL transgenes were used (∼50-60M cells used per replicate per condition). Cells were treated with pre-determined concentrations of doxycycline for 48h prior to experimentation. Plates of adhered cells were UV crosslinked in PBS on ice at an energy setting of 400 mJ/cm^2^. Immediately following crosslinking, cells were harvested, washed with ice cold PBS, centrifuged, and stored as cell pellets at −80°C.

Prior to performing pull-down, HaloLink resin was washed three times in PBST (PBS + 0.1% Tween-20) and blocked in 1% nuclease-free BSA (fraction V) for 20 minutes at room temperature. Resins were then washed three more times in PBST. Crosslinked cell pellets were incubated for 10 minutes on ice in lysis buffer (RIPA buffer (1M HEPES pH 7.4, 5M NaCl, 10% deoxycholate, 20% SDS, 10% NP-40), 1X Promega protease inhibitor cocktail, 1X TURBO DNase, 1x Mn^2+^/Ca^2+^ mix (1M MnCl_2_, 1M CaCl_2_), and Murine RNase inhibitor). Lysates were transferred to 1mL Bioruptor sonication tubes and sonicated on the “low setting” for 5 minutes (30s on/30s off) at 4°C. Following sonication, TURBO DNase and diluted RNase I (1:25 in PBS) were added to lysates and incubated at 37°C for 5 minutes with shaking at 1200 rpm. Immediately then murine RNase inhibitor was added to quench the reaction and lysates were then centrifuged at 15,000 rcf for 10 minutes at 4°C. ∼5% of the supernatant was reserved in −80°C to use later as an input control. The remaining lysate was added to the prepared HaloLink resin and incubated at 4°C overnight with rotation.

The next day, the following five CLAP wash buffers were prepared fresh and pre-warmed to 80°C prior to use: CLAP wash buffer 1 (20% N-lauroylsarcosine, 500mM EDTA, PBS); CLAP wash buffer 2 (1M HEPES pH 7.4, 5M NaCl, 10% NP-40, water); CLAP wash buffer 3 (1M HEPES pH7.4, 10% NP-40, 8M urea); CLAP wash buffer 4 (1M HEPES pH 7.4, 10% NP-40, 10% Tween-20, water); CLAP wash buffer 5 (1M HEPES pH 7.4, 10% NP-40, 500mM EDTA, water). Overnight captures were centrifuged and supernatants were discarded. Resins were washed twice with RIPA buffer. Resins were then washed three times each with each CLAP wash buffer in numerical order. For each wash, buffer was added to resins and incubated at 90°C for 3 minutes with shaking at 1200 rpm before centrifugation and removal of supernatant. Final resin pellets, as well as the reserved input samples, were then subjected to proteinase K elution by incubating with CLAP wash buffer 1 and proteinase K at 50°C for 20 minutes with shaking at 1200 rpm. The slurry was then transferred to Pierce micro-centrifuged columns to remove the resin. Supernatants were then used for RNA extraction with the Zymo RNA clean and concentrator kit. Eluted RNA was used for subsequent library preparation.

### CLAP library preparation

Input samples were first incubated with 1x TURBO DNase buffer and TURBO DNase at 37°C for 15 minutes with shaking at 1200 rpm and subsequently subjected to RNA purification again using the Zymo clean and concentrator kit. Eluted input RNA was then mixed with 1X FastAP buffer, fragmented at 91°C for 90s, and incubated with FastAP enzyme and murine RNase inhibitor for 10 minutes at 37°C with shaking at 1200rpm. At the same time, CLAP samples were mixed with 1X FastAP buffer, FastAP enzyme, TURBO DNase, and murine RNase inhibitor, and incubated for 10 minutes at 37°C with shaking at 1200rpm. Then, for all input and CLAP samples, PNK mix (1X PNK buffer, PNK enzyme, TURBO DNase) was added on top of the FastAP reaction and incubated for an additional 10 minutes at 37°C with shaking at 1200rpm. RNA was then purified using the Zymo RNA clean and concentrator kit.

To each sample, we then added the RIL19 adaptor (5’–/Phosphate/rArGrArUrCrGrGrArArGrArGrCrGrUrCrGrUrG/3ddC/ – 3’) in DMSO and incubated at 65°C for 2 minutes. Then ligation mix (1X T4 RNA ligase buffer, ATP, PEG8000, murine RNase inhibitor, and T4 RNA ligase high concentration) was added and incubated at room temperature for 1 hour and 15 minutes with rotation. Adaptor-ligated RNA was then cleaned with SILANE beads.

Eluted RNA was then incubated with reverse transcription adaptor AR17 (5’-ACACGACGCTCTTCCGA-3‘) at 65°C for 2 minutes. Reverse transcription mix (1X Maxima RT buffer, dNTPs, murine RNase inhibitor, Maxima RT enzyme) was added to the adaptor ligated RNA and incubated at 50°C for 20 minutes. Samples were then incubated with ExoI at 37°C for 15 minutes and quenched with EDTA on ice. To fragment RNA, the samples were incubated with NaOH at 80°C for 6 minutes and neutralized with equal amount of HCl. Reverse transcription product was then cleaned with SILANE beads. The resulting cDNA was then mixed with a splint adaptor (detailed in the published protocol) and 1X Instant Sticky Master Mix and incubated for 4 hours at room temperature with shaking at 1000 rpm. Each reaction was then cleaned with SILANE beads.

With the eluted cDNA samples we then performed PCR with NEBNext Universal primers for Illumina, NEBNext Index primers for Illumina, and 1X Q5 master mix. The following PCR conditions were used: (1X) 98°C 1 minute; (4X) 98°C 15s, 69°C 15s, 72°C 90s; (4X for inputs, 10X for IP) 98°C 15s, 72°C 90s; (1X) 72°C 2 min). PCR products then underwent 1.2X SPRI bead size selection and elution. Concentrations of the final barcoded libraries were measured on the Qubit and size measurements were calculated using the Agilent Tapestation prior to sequencing on a NextSeq2000.

### CLAP-seq data processing and analysis

The mapping of CLAP-sequencing was conducted according to published protocols from the Dr. Mitchell Guttman lab^35^. Adapters were trimmed from the paired end sequencing reads using cutadapt v4.9 from Trim Galore!. The RNA alignment was performed in two parts. First, the reads were aligned to hg38 genome reference containing both repetitive and structure RNAs using bowtie2. Second, the remaining reads were recycled and aligned to hg38 with STAR. After alignment, MarkDuplicates function from the Picard package was used to remove any PCR duplications. Finally, alignments from both mapping methods were finally merged into a single bam file for future analysis. CLAP peaks were identified by MACS2 peakcalling with IP and input from each condition using macs2 callpeak -t -c -f BAMPE -g hs -n -- nomodel --broad --broad-cutoff 1e-10 -p 1e-10 --keep-dup all. Peak distribution and associated genes were identified using HOMER module annotatePeaks.pl v4.11. NGS plots were generated use ngs.plot.r v2.61. For plotting, IP samples were normalized to their corresponding input using either signal subtraction or log2 fold change normalization with bamCompare/bigwigCompare. For signal quantification, read count files over peak regions were obtained from BAM files using featureCounts, followed by normalization to the sequencing depth.

### Western blot

Cells were collected, washed once in PBS, and boiled (95°C) in 1X SDS loading buffer (5X SDS loading buffer: 0.25M Tris-HCl (PH6.8), 0.5M DTT, 10% SDS, 50% glycerol, 0.25% BPB, 1% b-mercaptoethanol (b-ME) for 20 minutes. Lysates were run on homemade 10% SDS-PAGE gels in homemade 1X running buffer (2.5mM Tris Base, 19.2 mM Glycine, 0.01% SDS in H2O, pH 8.3). Protein was transferred to PVDF membrane (Millipore, 0.45 um) using freshly made homemade transfer buffer (1.2 mM Tris Base, 9.6 mM Glycine in H2O, pH 8.3) + 20% methanol on ice. After transferring, membranes were blocked with 5% blocking buffer in PBST (PBS + 0.1% Tween) for 45 minutes at room temperature. Membranes were then incubated with primary antibody suspended in 1% blocking buffer overnight at 4°C with rocking. The next day, membranes were washed three times for 10 minutes each wash with PBST. Membranes were then incubated with secondary antibody suspended in 1% blocking buffer for 1 hour at room temperature and subsequently washed three more times in PBST as before. Blots were covered with Pierce ECL Western Blotting Substrate according to the manufacturer’s instructions and imaged using film. Primary antibodies were diluted as follows: Flag (1:1000), MLLT1/ENL (1:1000), GAPDH (1:5000), CDK9 (1:1000), ELL2 (1:1000), LaminB1 (1:1000), a-Tubulin (1:1000). Secondary antibodies were used as follows: Anti-mouse or Anti-rabbit IgG HRP-linked (1:2500). For relevant quantifications, band intensities were measured in FIJI.

### Cycloheximide pulse chase

HEK293 cells expressing Flag-tagged ENL transgenes were incubated with doxycycline for 48h prior to experimentation. Cells were treated with 50 ug/mL cycloheximide for and collected at indicated timepoints. The intensity of each sample’s FLAG-ENL band was normalized to the intensity of that sample’s GAPDH. Intensities were measured in FIJI.

### Cell fractionation

Approximately 20M cells were harvested and washed with ice cold PBS. The cell pellet was flash freezed at −80°C and thawed on ice. Once pellets were thawed, 1mL of Buffer A (10mM KCl, 10mM HEPES pH 7.9, 1.5mM MgCl2, 0.5mM PMSF, 0.5mM DTT, protease inhibitors) was added and samples were incubated in a rotator for 20 minutes at 4°C. Samples were centrifuged at 3500 x g for 10 minutes at 4°C. Lysates were saved as the cytoplasmic fraction. The remaining pellet was incubated in an ∼equal volume (∼50uL) of Buffer C (20mM HEPES pH 7.9, 1.5mM MgCl2, 0.42M NaCl, 0.2mM EDTA pH 8, 25% glycerol, 0.5mM PMSF, 0.5mM DTT, protease inhibitors) in a rotator for 30 minutes at 4°C. Samples were centrifuged at 15,000 x g for 10 minutes at 4°C. Lysates were saved as the nuclear fraction. The remaining pellet was incubated in an ∼equal volume (∼50uL) of Nuclease inhibition buffer (150 mM HEPES pH 7.9, 1.5mM MgCl2, 150mM KOAc, 10% glycerol, 0.5mM PMSF, 0.5mM DTT, protease inhibitors) with RNase and DNase in a rotator for 30 minutes at 4°C. Samples were centrifuged at 12,000 rpm for 30 minutes at 4°C. Lysates were saved as the chromatin fraction. 30 ug of each fraction was loaded to a 10% SDS-PAGE gel and western blot was carried out as described above. For relevant quantifications, band intensities were measured in FIJI. LaminB1 and Tubulin serve as fractionation quality markers. Plotted for each fraction is the intensity of FLAG-T1 or FLAG-T1-3A relative to FLAG-T1.

### Immunoprecipitation

Approximately 20M cells were harvested and washed with ice cold PBS. The cell pellet was incubated in RIPA lysis buffer (15mM Tris-HCl pH 7.4, 1mM EDTA, 150mM NaCl, 1mM MgCl2, 10% glycerol, 0.1% NP-40, 0.5mM PMSF, 0.5mM DTT, protease inhibitors) for 20 minutes on ice. Samples were centrifuged at maximum speed for 10 minutes at 4°C. Lysates were reserved. Prior, 35 uL protein G beads per IP were washed three times with PBS + 0.01% Tween-20 and incubated with 5 ug FLAG antibody per IP for 5-6h in a rotator at 4°C. Beads were washed three more times with PBS + 0.01% Tween-20 before adding 900 uL of lysate. 50 uL of lysate was reserved for input controls in −80°C. Bead/lysate mixture was incubated overnight in a rotator at 4°C. The next day, beads/lysate mixtures were magnetized and washed three times with cold RIPA lysis buffer. 50 uL of 1X LDS loading buffer was added to the beads and samples were boiled for 10 minutes to elute proteins from beads. Beads were then magnetized and clarified lysate was saved for Western blot loading as described above. Input lysate was loaded at a volume equal to 5% of IP loaded. For relevant quantifications, band intensities were measured in FIJI. CDK9 and ELL2 serve as known ENL interaction partners. Plotted for each protein is its intensity in FLAG-T1 or FLAG-T1-3A IPs.

### Immunofluorescence staining and confocal microscopy

Cells (150K/well) were seeded on 24-well plates containing coverslips and incubated 24-48h. Cells were fixed with 4% PFA in PBS for 10 min at room temperature, washed with PBST (PBS + 0.1% Tween-20) three times (5 min/wash), and then permeabilized with PBS containing 0.1% Triton-X 100 for 10 min, and washed three times with PBST as before. Subsequently, cells were blocked in PBS with 10% goat serum for 30 min at room temperature. Cells were then incubated overnight at 4°C with primary antibodies (1:300) in 1% BSA in PBS. The next day, cells were washed three times in PBST, incubated with secondary Alexa Fluor-conjugated antibodies (1:250) in 1% BSA for 1h at room temperature, and washed three more times in PBST. The coverslips were then mounted face down on slides in DAPI-containing mounting medium. Primary antibodies used: Flag (Sigma F1804) and MLLT1/ENL (Millipore ABE2596). Secondary antibodies used: goat anti-Rabbit or anti-Mouse IgG (H+L) conjugated with Alexa Fluor 488, 568, or 647 (Invitrogen). Images were captured on Zeiss LSM880 or Leica Stellaris 5 confocal microscopes.

### DNA-FISH with immunofluorescence staining

Poly-L-lysine was first added to wells of rubber gaskets adhered to glass microscopy slides and removed/dried prior to cell plating. Cells were then plated in the gasket wells and allowed to adhere at 37°C for 1.5-2h (HEK293 and MDA-MB-231cells) or 20 minutes (LSK cells). Slides were then fixed in 4% PFA for 10 minutes at room temperature in coplin jars followed by three 10 minute washes in room temperature (RT) PBS. **DNA-FISH Part 1**: Slides were permeabilized in 0.5% Triton-X-100 in PBS for 15 minutes at RT and washed once in PBS for 5 minutes at RT. Next slides were incubated in 70% EtOH, 90% EtOH, and 100% EtOH for 2 minutes each time. Slides were then washed once in 2X SSCT (1X SSC + 0.1% Tween-20) for 5 minutes at RT and incubated once in 70% FMM buffer (70% formamide, 1X SSC, water) for 5 minutes at RT. Slides were then transferred to 70% FMM buffer pre-heated to 37°C for 1 h. Then primary FISH probe hybridization solution (50pmol primary probe, 100uM dNTP, 50% FMM, 1X dextran-sulfate PVSA hybridization mix, 10ug RNase A, water) was added to each slide, sealed with a coverslip and rubber cement, and denatured on a water-immersed heat block for 30 minutes at 83°C. Slides were then transferred to a humidity chamber and incubated overnight at 37°C. **DNA-FISH Part 2:** The next day, slides were washed once in 2X SSCT for 10 minutes at 60°C, once in 2X SSCT for 10 minutes at RT, and once in 0.2X SSC for 10 minutes at RT. Slides were then incubated with secondary FISH probe for 2 h at RT protected from light under coverslips sealed with rubber cement (secondary probe mix: 10pmol secondary probe, 10% FMM, 1X dextran-sulfate PVSA hybridization mix, water). Slides were then washed again in 2X SSCT 60°C, 2X SSCT RT, and 0.2X SSC as before. **IF Part 1:** Slides were next fixed again in 2% PFA for 10 minutes at RT, washed once in PBS for 5 minutes at RT, permeabilized with 0.1% Triton-X-100 for 15 minutes at RT, and washed three times in PBST (PBS + 0.1% Tween-20) for 10 minutes each at RT. Slides were then blocked in 1% BSA in PBST for 45 minutes at RT. Then primary antibody solution (1:300 primary antibody in 1% BSA) was added to each slide, sealed with a coverslip and rubber cement, and incubated overnight at 4°C in a humidity chamber. **IF Part 2:** The next day, slides were washed three times in PBST for 10 minutes each time. Then secondary antibody solution (1:250 secondary antibody in 1% BSA) was added to each slide, sealed with a coverslip and rubber cement, and incubated for 2 h at RT. Slides were washed again three times in PBST for 10 minutes each time. Finally, slides were incubated in DAPI staining buffer (2X SSC + 1:10,000 Hoescht stain) for 5 minutes at RT and washed once in PBS for 5 minutes at RT. Coverslips were added to each slide with anti-fade mounting media and sealed with clear nail polish.

Primary probes used: Probes were designed to cover ∼100kb regions (4 40mer probes per kb) of the human *HOXA9-13* locus, human *CBX3* locus, human *NR2F2* locus, human *CHD2* locus, mouse *Hoxa9-10* locus, or mouse *Meis1* locus. Secondary probes were dye conjugated. Primary antibodies used: Flag for HEK293 and MDA-MB-231 cells (Sigma F1804) and MLLT1/ENL (Millipore ABE2596) for LSK cells. Secondary antibodies used: goat anti-Rabbit or anti-Mouse IgG (H+L) conjugated with Alexa Fluor 488, 568, or 647 (Invitrogen). Images were captured on Zeiss LSM880 or Leica Stellaris 5 confocal microscopes.

### ChIP-qPCR and ChIP-sequencing

Between 20-25M cells were used per ChIP. Cells were collected from plates and fixed in freshly made 1% PFA in PBS (1mL 1% PFA per 10M cells). For Flag ChIP, fixation was performed for 10 minutes at room temperature with intermittent inversion. PFA was quenched with 125mM Glycine for 5 minutes at room temperature and the cell pellet was then washed three times with PBS. Cells were then resuspended in RIPA3.0 buffer (0.1% SDS, 1% Triton-X-100, 10mM Tris-HCl pH7.4, 1mM EDTA pH8, 0.1% NaDOC, 0.3M NaCl) freshly supplemented with 1/40 volume of 10% sarkosyl, 1mM DTT, and protease inhibitor. For MDA-MB-231 ChIPs, an extra 0.1% SDS (final 0.2% total) was added to the sonication buffer to improve sonication. Cells were sonicated on a Covaris Ultrasonicator. Concurrently, primary antibody was incubated with Protein A Dynabeads and then washed three times with PBS + 0.1% Tween-20. A small volume of sonicated lysate was reserved as an input control. Remaining lysate was rotated overnight at 4°C with the primary antibody-bound Dynabeads. For MDA-MB-231 ChIPs, sonicated lysates (which contain extra SDS in the sonication buffer) were diluted with non-SDS containing sonication buffer to a final concentration of 0.1% SDS before adding to the antibody-bound beads. While extra SDS improved sonication, this SDS can interfere with IP and must be re-diluted before IP. The next day, beads were washed three times with low salt buffer (150mM NaCl, 0.1% SDS, 1% Triton-X-100, 1mM EDTA pH8, 50mM Tris-HCl pH8), three times with high salt buffer (500mM NaCl, 0.1% SDS, 1% Triton-X-100, 1mM EDTA pH8, 50mM Tris-HCl pH8), and three times with LiCl buffer (150mM LiCl, 0.5% NaDOC, 0.1% SDS, 1% NP-40, 1mM EDTA pH8, 50mM Tris-HCl). Beads were then washed once with TE buffer (1mM EDTA, 10mM Tris-HCl pH8) + 50mM NaCl. Elution buffer (1% SDS, 50mM Tris-HCl ph8, 10mM EDTA, 200mM NaCl) was then added to the beads and incubated at 65°C with shaking for 0.5h. Beads were then centrifuged and supernatant was transferred to a new tube and incubated overnight at 65°C. At this point elution buffer was also added to reserved input lysate and incubated overnight at 65°C. The next day, the samples were first incubated with RNase A for 1h and 37°C and then with Proteinase K for 1h at 55°C. DNA was purified using the Qiagen PCR purification kit and eluted in PCR water.

For qPCR, ChIP DNA was diluted 50x in water and qPCR was performed using SYBR green master mix. Primers used are listed in Supplementary Table 11. For ChIP-sequencing, ChIP DNA was subjected to library preparation using the Vazyme Universal DNA Library Prep Kit for Illumina. Prior to sequencing, libraries were quantified and size confirmed using the Qubit and Tapestation according to the manufacturers’ instructions. Final libraries were sequenced on an Illumina Nextseq 2000.

Reads were aligned to the human reference genome (hg19) by the package Bowtie2 v2.5.4 with the default parameters. The package samtools v1.9 was used for converting to BAM format, sort-ing, removing duplicates, and indexing. In ChIP-seq analysis, we utilized the corresponding input data as a control during peak calling. The chromatin distribution of the peaks was determined by HOMER module annotatePeaks.pl v 4.11. Peaks are listed in Supplementary Tables 2, 4). To make the heatmaps, the filtered, sorted, and indexed BAM files were converted to bigwig files using the bamCoverage from deeptools v3.5.6 and then normalized by counts per million (CPM). Heatmaps were generated using the module computeMatrix and plotHeatmap from deeptools v3.5.6.

### RNA extraction, cDNA synthesis, and RT-qPCR

Total RNA was extracted using Qiagen RNeasy kits according to the manufacturer’s instructions. Reverse transcription was performed using the High-Capacity cDNA Reverse Transcription Kit (Applied Biosystems) according to the manufacturer’s protocol. Quantitative real-time PCR was performed using SYBR green master mix. Primers used are listed in Supplementary Table 11.

### RNA-sequencing and Gene Set Enrichment Analysis (GSEA)

RNA was extracted as described above. Library preparation was performed with 500 ng total RNA beginning with the NEB polyA mRNA magnetic isolation module followed by NEB library preparation kit, both according to the manufacturer’s instructions. Prior to sequencing, libraries were quantified and size confirmed using the Qubit and Tapestation according to the manufacturers’ instructions. Final libraries were sequenced on an Illumina Nextseq 2000.

Reads were aligned to the mouse reference genome (mm10) using HISAT2 v 2.2.1 with default parameters. The package featureCounts v2.0.2 was used for counting the mapped reads in each mouse gene. Transcript per million (TPM) was used for normalizing the gene expression. Differentially expressed genes between conditions were statistically determined by R package DESeq2 v1.38.3. Genes with adjusted *P* value < 0.05 and fold change ≥ 1.5 were reported as differential genes. Differentially expressed genes in LSK are listed in Supplementary Table 9.

For gene set enrichment analysis, custom gene signatures were made from our ChIP-sequencing datasets (Supplementary Tables 5, 6, 8). Briefly, normalized read counts for ENL-T1 or -T2 peaks were ranked and the inflection point determined each peak to be top or low-bound. Top-bound peaks were annotated with a corresponding gene and this list of top-bound genes was used as the gene signature. DESeq2 files used in the analysis were pre-ranked. GSEA was performed with the GSEA_4.3.2 desktop application.

### RNA-FISH with Halo staining

HEK293 cells expressing Halo-tagged ENL transgenes were seeded on poly-L-lysine coated coverslips and grown for 48h in the presence of doxycycline. Live cells were then incubated with 250 nM JF657 Halo ligand for 30 minutes and washed five times in pre-warmed media (5 minutes per wash). Cells were then washed once in PBS + 1% RVC (ribonucleoside vanadyl complex), fixed in 4% PFA + 1% RVC for 10 minutes at room temperature, and then washed twice more in PBS 1% RVC. The cells were then permeabilized in 70% ethanol and 1% SDS overnight at 4°C. The next day, the ethanol/SDS solution was removed and the cells were washed twice with 2xSSCT (2xSSC + 0.1% Tween-20) + 1% RVC at room temperature for 5 minutes per wash. Primary probe hybridization mix (2x DSM (dextran sulfate PVSA), 50% formamide, 1% SDS, 30 nM primary probe, 1% RVC) was assembled and added dropwise onto a parafilm sheet sitting on top of a damp paper towel in a petri dish. Coverslips were lifted and placed cell-side down onto the hybridization mix drops and denatured at 70°C for 4 minutes. The apparatus was then incubated at 42°C overnight. The following day, the coverslips were transferred back to 24-well plates and washed twice with 2xSCCT + 1% RVC at room temperature, four times with 2xSCCT + 1% RVC at 65°C, and twice more with 2xSCCT + 1% RVC at room temperature. Each wash was 5 minutes. Secondary probe hybridization mix (2x DSM (dextran sulfate PVSA), 50% formamide, 1% SDS, 50 nM dye-conjugated secondary probe, 1% RVC) was assembled and added dropwise onto a parafilm sheet sitting on top of a damp paper towel in a petri dish. Coverslips were lifted and placed cell-side down onto the hybridization mix drops and incubated for 1h at 37°C. Following secondary hybridization, the coverslips were transferred back to 24-well plates and washed four times with 2xSCCT + 1% RVC at 37°C. The coverslips were finally mounted cell side down on slides in DAPI-containing mounting medium. Images were captured on a Zeiss LSM880 confocal microscope.

Primary probe used: Probes were designed to cover a ∼750bp region (19 60mers) of the human *HOXA13* intron. Secondary probes were dye conjugated.

### Murine hematopoietic stem cell harvest and culture

Whole bone marrow was extracted from euthanized C57BL/6 mice using lower limb centrifugation and subjected to RBC lysis. cKit+ cells were then isolated using the EasySep Mouse CD117 (cKIT) Positive Selection Kit (StemCell). Cells were then stained with Zombie Aqua live/dead dye and cell surface markers (hematopoietic lineage markers, Sca-1, cKit) prior to sorting out Lin^-^Sca-1^+^cKit^+^ cells on a BD FACSAria. Lineage markers used are CD3, CD19, CD11b, B220, TER119, Gr-1, NK1.1, CD11c, TCRβ, and TCRγ/δ. LSK cells were cultured for 48h prior to subsequent virus transduction. Cells are cultured in IMDM + GlutaMAX supplemented with 10% FBS, 100 U/mL penicillin-streptomycin, 20 ng/mL mFlt3L, 20 ng/mL mIL6, 100 ng/mL mSCF, 20 ng/mL mTPO, 1:1000 0.1M betamercaptoethanol, 1:100 200mM L-glutamine.

### Murine hematopoietic stem cell viral transduction

For LSK cell infection, the day prior to infection 420 uL of RetroNectin reagent (60 ug/mL) was added to untreated 24-well cell culture plates and incubated overnight at 4°C. The next day, RetroNectin was removed and wells were blocked with PBS + 2% BSA for 30 minutes at room temperature. Wells were then washed once with Hank’s balanced salt solution (HBSS). After removing the HBSS, desired volume of retrovirus was added to the wells. Plates were centrifuged at 4°C 3000 rpm for 1h. Next, retroviral supernatant was removed and cells were added to the virus-loaded wells with 10 ug/mL polybrene and spinfection was carried out at 37°C 1500rpm for 1.5h. Plates were then incubated overnight. Cell sorting was performed 48-72 hours post-infection.

### Colony formation assay

6000 cells were resuspended in 600 uL of IMDM and added to 3mL of MethoCult methylcellulose media supplemented with 10 ng/mL mIL3, 10 ng/mL mIL6, and 20 ng/mL mSCF. 1 mL of the mixture was then added to each well of a 6-well plate (3 replicates total, 2000 cells per replicate). Plates were placed in the incubator and colonies were counted ∼1-1.5 weeks after plating. Subsequent re-platings were carried out in the same way.

### Bone marrow transplantation and engraftment measurements

LSK cells (CD45.2) were isolated and transduced with ENL transgenes as described above. Transduced LSK were mixed at a 1:3 ratio with RBC-lysed whole bone marrow from a CD45.1/2 mouse (100K LSK: 300K support cells). 400K cells resuspended in 100 uL PBS were injected retro-orbitally to each CD45.1 recipient mouse (B6.SJL-*Ptprc^a^ Pepc^b^*/BoyJ, Jackson Labs #002014). The morning and afternoon prior to injection, CD45.1 recipient mice underwent 2 rounds of lethal irradiation (5.5 Gy per round, 11 Gy total).

Beginning 2 weeks after transplantation, engraftment was measured by peripheral blood staining. Peripheral blood (PB) was harvested by submandibular bleeding into EDTA coated tubes. ∼15 uL PB was lysed in 1X RBC lysis buffer for 6 minutes on ice, quenched with FACS buffer (PBS + 2% FBS), and washed once in FACS buffer. Cells were stained with CD45.1 and CD45.2 antibodies (1:200) in FACS buffer for 30 minutes on ice protected from light. Cells were washed once more in FACS buffer and resuspended in FACS buffer + 1:2000 SYTOX blue dead cell stain. Samples were then run on an Attune NxT flow cytometer. Engraftment was measured approximately every 2 weeks for 1 year following transplantation.

### Statistics and reproducibility

All experimental observations were confirmed by independent repeat experiments. Experimental data is presented as mean ± standard deviation unless stated otherwise. Statistical significance was calculated by two-tailed, unpaired or paired Student’s t-test on two experimental conditions with *P* < 0.05 considered statistically significant unless stated otherwise. The number of replicates and the statistical test used were indicated in the corresponding figure legends. Statistics were performed using GraphPad Prism 10 and RStudio. No data were excluded from the analyses. No statistical methods were used to predetermine sample size. The experiments were not randomized. The investigators were not blinded to allocation during experiments and outcome assessment.

